# Gut Stem Cell Secretome Therapy Promotes Durable Neuroprotection and Functional Recovery After Stroke

**DOI:** 10.64898/2026.11.03.763998

**Authors:** Kathiresh Kumar Mani, Ann M. Alex, Kaylin Pickle, Zara Akbari, Akruthi Obhrai, Asmitha Senthil Kavithil, Netra Ashokraj, Srihari Ravindra Jadhav, Emily Stone, Nahla Adlouni, Gowtham Saksham, Kate A. Miller, Saarah Malik, Farida Sohrabji

## Abstract

**Background:** Stroke causes rapid neuronal death and gut barrier disruption in the acute phase, with persistent cognitive-affective impairment in the chronic phase. We previously showed that transplants of intestinal epithelial stem cells ameliorated the acute and chronic impairment of stroke. The current study tested the efficacy of a cell free strategy, specifically, the secretome of intestinal epithelial stem cells.

**Methods:** Middle-aged male and female rats underwent middle cerebral artery occlusion or sham surgery and received intestinal epithelial stem cell secretome from healthy sex-matched adult or control media. Behavioral assays (social interaction, novel object recognition and Barnes maze), gut histology, markers of gut permeability, serum cytokines were performed at acute and chronic time points. Group differences were assessed by two-way ANOVA.

**Results:** In the acute phase, vehicle-treated stroke animals displayed histopathological changes in the gut including villus blunting, loss of goblet cells, increased markers of gut permeability and elevated systemic inflammation. In the chronic phase, stroke resulted in cognitive/affective changes including social withdrawal, recognition memory and spatial memory deficits. In middle-aged males, three doses of IESC secretome treatment 4h/24h/48h after stroke restored mucosal architecture, goblet cells and dampened inflammatory cytokines in the acute phase. Cognitive and social outcomes, tested 4+ weeks after stroke, were significantly improved compared to vehicle-treated stroke animals. However, in middle-aged females treated with the same regimen of intestinal epithelial stem cell secretome, gut architecture was not restored and inflammatory cytokines remained elevated. Long term cognitive impairment was only partially improved in this group. Proteomic analysis of the secretome revealed a largely shared core of proteins between males and females with sex differences in pathway level regulation. Subsequently, another cohort of middle-aged females was administered four doses of secretome 4h/12h/24h/36h after stroke. This regimen restored gut architecture and dampened inflammatory cytokines in the acute phase, and improved long term cognitive outcomes.

**Conclusions:** Collectively, these studies underscore the therapeutic potential of IESC secretome and the importance of tailoring post stroke therapy by demographics such as biological sex. These data further support the hypothesis that gut ‘repair’ is tightly associated with long-term improvement in stroke outcomes.

## INTRODUCTION

Stroke is the second leading cause of mortality and a major contributor to chronic disability (1–3). Cognitive deficits are exceedingly common after stroke, nearly 70% of survivors develop some degree of cognitive impairment within the first year, and a substantial proportion progress to post-stroke dementia (4–9). Therapeutic options for stroke remain strikingly limited, either by a narrow safe treatment window as in the case of thrombolytics or requiring specialized resources and strict patient selection, as in endovascular thrombectomy, leaving the majority of individuals without effective acute intervention (10).

Growing evidence indicates that stroke is ‘whole body’ problem, and implicates the gastrointestinal system as a critical, yet underappreciated, modulator of stroke pathophysiology. Following stroke, nearly half of all patients experience complications such as gastrointestinal bleeding, ileus, fecal incontinence, or dysphagia, conditions that are linked to poorer neurological recovery and higher mortality (11–14). Experimental models further demonstrate that stroke compromises intestinal barrier integrity and induces marked alterations in gut microbial composition (15, 16), driving secondary injury through inflammatory, neural, and metabolic pathways (17).

Our previous studies have shown that middle cerebral artery occlusion (MCAo) disrupts intestinal architecture and barrier function rapidly after injury, accompanied by sustained elevations in inflammatory cytokines such as IL-17A that correlate with the emergence of affective disturbances and cognitive decline weeks later (18, 19). Consequently, we leveraged the linked observations of stroke-induced intestinal dysmorphology in the acute phase and stroke-induced cognitive-affective impairment in the chronic phase, to focus on therapies that repair the gut. Our recent work showed that transplantation of intestinal epithelial stem cells (IESCs) shortly after stroke ameliorated pathological gut features and improved neurological functional outcomes in middle-aged male and female animals (18). While these findings are promising, direct stem cell–based therapy in clinical settings face substantial logistical and biological barriers, including immunological rejection, tumorigenicity, and ethical concerns (20, 21). Based on the evidence from preclinical studies that the regenerative and neuroprotective actions of transplanted stem cells are largely mediated by their secreted paracrine factors (22, 23), the present study evaluated the therapeutic potential of an IESC-derived secretome. Our data demonstrate that IESC-secretome improves gut barrier integrity and concomitantly enhances acute stroke outcomes in both sexes. Surprisingly, chronic stroke outcomes showed sex specific responses to secretome treatment, where females required additional/early dosing of IESCs-secretome to achieve comparable benefits as males. As in our previous studies, we identified that improved cognitive outcomes were associated with improved gut histology and decreased levels of inflammatory cytokines. Our data illustrates the importance of precision, sex-informed dosing strategies of secretome-based therapeutics.

## METHODOLOGY

### Animal model

This study utilized adult (5-7m) and middle-aged (9-11m) male and female Sprague Dawley rats, purchased from Inotiv Laboratories (IN). Rats were maintained in a constant 12-h dark: 12-h light cycle with ad libitum food and water. All animals were fed pelleted food (Harlan 8604 Teklad diet) for at least 4 weeks prior to their assignment to the study. All procedures were reviewed and approved by the Texas A&M University Institutional Animal Care and Use Committee (IACUC 2023-0307) in accordance with OLAW guidelines for the humane treatment of animals in research. Daily vaginal smears were obtained to confirm that middle-aged females were acyclic (24). In all cases, middle aged animals were hosts; adult animals were donors (Fig.S1-Experimental strategy).

### Preparation of IESCs-secretome from donors

Primary IESCs were isolated from adult male and female rats using procedures described in our previous publication (18). IESC were cultured in organoid basal medium containing 100X L-glutamine (Sigma-Aldrich, Catalog G7513, 20ul/ml), 1M HEPES (Invitrogen, Catalog 15630–056, 20ul/ml), 50X B27-supplement (Invitrogen, Catalog 17504–044, 40ul/ml) and 100X N2-supplement (Invitrogen, Catalog 175024–048, 20ul/ml), growth factors (1000x N-acetylcysteine (Sigma-Aldrich, A9165, 1 mM), 1000x murine EGF (Peprotech, 315–09, 50 µg/ml), 500x murine Noggin (Peprotech, 250–38, 50 µg/ml), 200x murine R-spondin 1 (R&D Systems, 3474, 100 µg/ml), 10 mM/ml of A8301 (Sigma-aldrich 70024–90-7) and 10 mM/ ml Y-27632 (ROCK inhibitor, Stem cell Technologies, 72304). Once cultures reached a count of ∼500 organoids (on day 3), the medium was aspirated and cells were washed twice with Dulbecco’s phosphate buffer saline (DPBS). Cultured plates were then refed with plain DMEM medium without FBS and cultures were incubated for another 24hr. The media was then collected at 24hr and centrifuged at 3000rpm for 15min at 4°C to remove the cell debris. The supernatant was then filtered through a sterile filter with a pore size of 0.22 µm to remove any remaining particles and ensure sterility. Finally, IESC-secretome was stored immediately at −80°C until further use.

### Proteomics analysis of IESC-secretome

IESC secretome was analyzed by LC-MS/MS using an Ultimate 3000 nano-LC system coupled to an Orbitrap Fusion tribrid mass spectrometer (Thermo Scientific) at the Texas A&M Protein Core. Proteomic analyses were performed on 27 samples: basal media controls (BM; n = 9), female secretome (conditioned media) from adult female rat IESC culture (IESC-female secretome; n = 9), and male secretome (conditioned media) from young rat male IESC (IESC-male secretome; n = 9).

#### (i) Protein quality filtering and background subtraction

Raw MS/MS spectra of peptide ions were subsequently processed using Proteome Discoverer 2.4 software (ThermoFisher Scientific) against a Rattus norvegicus protein database obtained from GenBank (www.ncbi.nlm.nih.gov). Proteins were retained only if they were supported by at least two unique peptides and were classified as high confidence (≥2 unique peptides; exclusion of medium-confidence accessions) in Proteome Discoverer; two medium-confidence proteins were excluded at this step. For the BM group, we defined the background as the union of all proteins detected in any of the nine BM replicates (n = 23), ensuring that all potential media-derived contaminants were flagged for removal.

For IESC-secretome samples, we identified proteins that were consistently detected across all nine replicates within each sex (females: n = 958; males: n = 997) and across all 18 IESC-secretome samples combined (n = 860). Individual female IESC secretome samples contained 1,254-1,491 proteins, and male IESC secretome samples contained 1,269-1,461 proteins. Before BM subtraction, 860 proteins were shared between male and female IESC-secretome and IESC-secretome and were present in every IESC-secretome sample. To isolate IESC-secretome-specific proteins, all proteins detected in any BM replicate (23 proteins) were removed from both male and female IESC-secretome datasets. We first assessed the consistency of BM protein detection: only 2 of 23 BM proteins were observed in all nine BM samples, whereas the remaining 21 appeared in a subset of replicates. We then queried each BM protein across the 18 individual IESC-secretome samples: 8 proteins were present in all CM samples, 2 were detected in 17 of 18 samples, and 12 were found in four or fewer IESC-secretome samples, including 6 that were never detected in IESC-secretome. After BM subtraction, the final IESC-secretome datasets comprised 949 proteins in females and 852 in males of which 97 were unique to female IESC secretome, 136 were unique to male IESC secretome, and 852 were shared between sexes

#### (ii) Ingenuity Pathway Analysis and rationale

BM-subtracted IESC-secretome protein sets were submitted to Ingenuity Pathway Analysis (IPA) (QIAGEN Inc., https://digitalinsights.qiagen.com/IPA) using the Core Analysis workflow (Kramer et al., 2014) with the species filter set to Rattus norvegicus. We provided, for each protein, both its accession identifier and associated expression statistics (log₂ fold change and Benjamini-Hochberg adjusted p-value). Of the 988 proteins submitted in the male dataset, 954 mapped to the IPA knowledge base, and of the 949 proteins in the female dataset, 913 were mapped. We filtered canonical pathway results to include only those with an absolute activation z-score ≥ 2 and a Fisher’s exact test p-value < 0.05. These combined thresholds enrich pathways that are both significantly overrepresented in the IESC-CM datasets and have a robust predicted direction of regulation.

#### Quantification of Growth Factors in IESC Secretome

Although mass spectrometry provides broad protein profiling, small and low abundance signaling molecules such as growth factors are often under-represented due to sample processing losses, ionization inefficiency, and competition with highly abundant peptides. Accordingly, a membrane-based antibody array was used to detect 14 rat growth factors from liquid samples (RayBiotech, GA, Cat # AAR-GF-1-2). Growth factors that were detected on this array were the assayed by quantitative ELISA kits specific for each: VEGF (MyBioSource 724516), PDGF-A (ThermoFisher), BDNF (ThermoFisher), and IGF-1 (R&D Systems).

### Preparation of Hosts

Rats designated as hosts were randomly assigned to the following groups: Sham (no MCAo), MCAo with vehicle (Basal media), or MCAo with IESCs-secretome. In all, 35 middle aged females (7 sham, 16 MCAo/Vehicle and 12 IESC-secretome), 37 middle aged males (7 sham, 16 MCAo and 14 IESC-secretome) were used for the acute study (3 doses). For the chronic study, 37 middle aged females (9 sham, 14 MCAo/Vehicle and 14 MCAo/ IESC-secretome) and 36 middle aged males (8 sham, 15 MCAo and 13 IESC-secretome) were used (3 doses). In the 4 doses study, 28 middle-aged females (7 sham, 11 MCAo/Vehicle and 10 MCAo/IESC-secretome) and 29 middle-aged females (7 sham, 11 MCAo/Vehicle and 11 MCAo/ IESC-secretome) were used for the acute and chronic study respectively.

### Middle Cerebral Artery Occlusion (MCAo)

Animals were subjected to stereotaxic surgery to occlude the left middle cerebral artery as previously reported (25). Briefly, endothelin-1 (3 µl, 1:2 dilution in DPBS of 1 mg/ml endothelin-1 stock; American Peptide Company, Inc., CA) was injected at the following stereotaxic coordinates relative to bregma: + 0.9 mm anterior/posterior, + 3.4 mm medial/lateral, and −8.5 mm relative to the dura. The same procedure was followed for sham surgeries without the injection of endothelin-1. Stroke-induced mortality was recorded in all experiments.

### IESCs-secretome administration

Dosing regimen: The three-dose IESC-secretome treatment was administered intraperitoneally (i.p.) (i.p.) 4, 24 and 48h (0.5 mL per dose) after MCAo to match the dosing regimen used for IESC cell transplants (18). Subsequently, a four dose treatment was administered i.p. to a separate cohort of females at 4,12,24 and 36h after MCAo (0.5 mL per dose). For all experiments, each recipient received secretome derived from a unique donor (adult, 5–7 months old) of the same sex.. The timeline (experimental strategy) of the acute and chronic experiments is shown schematically in Fig. S1.

### Acute stroke outcomes

#### Infarct volume

Infarct volume analysis was determined using our previous procedures (18, 19). Total brain infarct was calculated from 3 slices per animal and expressed as the ratio of infarct volume in the ischemic hemisphere to the total volume of the non-ischemic hemisphere.

#### Behavioral analysis

All testing was performed between 9 am and 1 pm to minimize the contribution of circadian variations to behavioral performance as described previously (18, 19, 26). All tests were performed and scored by experimenters who were blind to the treatment condition. For assessing sensorimotor function, the adhesive tape removal test (ART) and the vibrissae evoked forelimb placement task (VIB) were performed before (Pre, −2 days) and after (2 and 4 days) stroke (18, 19). A separate set of animals were prepared for cognitive-affective behavioral analysis which were performed 4 weeks after stroke. The sequence of behavioral testing was Social Interaction (SI), Novel Object Recognition Task (NORT) and Barnes maze test. In general, subsets of all the three groups (Sham/MCAo+Vehicle/MCAo+ IESC-Secretome) of animals were assessed on the same day for a particular behavioral test. For most assays, tests were performed pre and post stroke. Due to multiple behavior tests, animals were not randomized by baseline values but were compared pre and post using repeated measures analysis.

##### Social interaction test

A three-chambered Plexiglass box was used to assess social interaction (18, 27). Time spent in the ‘social chamber’ which contained a conspecific same sex rat was recorded.

##### Tests of cognitive function

Two tests were performed. Novel object recognition task (NORT): Object recognition memory was tested as described previously (18, 27, 28). The discrimination index was calculated as follows: DI: [time spent with novel object] -[time spent with familiar object]/[time spent with novel object] + [time spent with familiar object].

##### Barnes Maze

This test was performed to assess spatial learning and recall as described previously (18, 27, 28) Latency to find the escape box as well as velocity and distance traveled during each trial was recorded. Following spatial acquisition, a probe test was conducted to assess spatial reference memory. The Ethovision software (Noldus) was used to analyze measures during the learning phase and time spent in the “target quadrant” during the recall portion of the test.

### Enzyme-linked immunosorbent assay

Serum was collected via saphenous draw at baseline (pre-stroke), 4 days and 30 days post-stroke. ELISA assays were used to determine Lipopolysaccharide (LPS) (Mybiosource, MBS268498) and Intestinal Fatty Acid-Binding Protein (iFABP) (Mybiosource, MBS3807789) as a measure of gut permeability. GFAP (Glial Fibrillary Acidic Protein) assay (MyBioSource, MBS2505953) was used to assess blood-brain barrier integrity during the acute phase(4dp). Serum cytokine/chemokine were quantified using a rat cytokine/chemokine multianalyte panel (Millipore, MA). All procedures were performed according to the manufacturer’s directions and our published procedures (16, 18, 19).

### Gut Histology

Gut health was assessed by a set of measures including routine histology (H&E staining), Periodic Acid Shiff staining, and surrogate markers of gut permeability. Gut histology focused on the distal ileum, a region where severe morphological changes are seen after stroke (16, 18). Cryosections (10µm) were collected on glass slides (3 per slide) and analyzed for hematoxylin and eosin (H&E) staining and Periodic acid-Schiff (PAS) as previously described (18, 19). Sections were imaged with the Olympus VS120 virtual slide scanning system. Analysis of H&E and PAS staining: In the H & E images, the villus length to villus width ratio was assessed for every villus on three sections per animal using the ImageJ software (16, 18). The length of the villus was measured from its base at the crypt to its tip and its width was measured at its widest point. The number of crypts at the base of each villus was also counted. In PAS-stained images, the number of goblet cells was counted for each villus on 3 sections per animal. Goblet cells were identified by their characteristic shape, pink coloration and location along the borders of the villi.

### Magnetic resonance imaging

Brain imaging was performed on postmortem tissue. Male and female rats were imaged on a Siemens 3 tesla Magnetom Verio Magnetic resonance imaging (MRI) scanner at the Texas A&M Institute for Preclinical Studies using a T2 weighted (T2WI) imaging sequences with a 32-channel body coil. Imaging parameters were = 5760/27 ms, matrix = 208 × 256, and slice thickness at 1 mm. T2WI sequence parameters were Repetition Time/Time to Echo (TR/TE) = 2830/10.6 ms, matrix = 256 × 256, and a slice thickness of 0.75 mm. Images were processed and analyzed using OsiriX DICOM viewer and Fiji (for Image J2) to delineate the volumes of hyperintense lesions and brain hemispheres. For each slice, the measured area was multiplied by the slice thickness, all individual slices were summed to obtain the total hyperintensity volume.

### Statistical Analysis

Data were primarily analyzed using parametric statistical tests (Prism Graphpad). Survival plots were calculated using the Kaplan Meier test. For behavioral tests, pre-and post-stroke scores were analyzed by two-way ANOVA for repeated measures and Tukey’s test for post hoc analysis. For all other comparisons, an unpaired Student’s *t* test or a one-way ANOVA were used. Group differences were considered significant at p≤0.05 in each case. All data are expressed as mean ± S.E.M. Specific animal numbers used for an assay are described in each figure legend.

#### Proteome analysis

Protein filtering and downstream data handling were performed in R (v4.4.2) using the tidyverse suite (v2.0.0; (Wickham et al., 2019)). Canonical pathway results from IPA were visualized using ggplot2 (v4.0.2;(Wickham, 2016)), patchwork (v1.3.2; (Pedersen, 2025)), and scales (v1.4.0;(Wickham et al., 2025). To characterize the overall effect of secretome treatment on gut barrier and immune biomarker profiles, principal component analysis (PCA) was performed separately for the acute and chronic datasets using the R package FactoMineR (v2.13;( Lê et al., 2008)). Variables were mean-centered and scaled to unit variance and only animals with complete data across all variables were included in each analysis. Animal scores on the first two principal components were plotted using ggplot2, colored by treatment group. For the acute (4dp) dataset, six biomarkers were included: villus length-to-width ratio, goblet cell count, IL-17A, IL-6, IL-10, and TNF-α. For the chronic (30d) dataset, behavioral variables were plotted, using delta values (post-minus pre-treatment) for social interaction and novel object recognition testing, and latency on the terminal training day (Day 4) of the Barnes maze test.

## RESULTS

Our previous study using IESC transplants showed that long term improvement in cognitive function was associated with early improvement in gut permeability and histomorphology. Thus, we first assessed whether IESC secretome would have a similar reparative effect on the gut in the acute phase.

### IESCs-secretome treatment exerts a sex-specific effect on stroke-induced disruption of gut architecture in middle-aged rats

H&E-stained sections of the ileum from Sham, MCAo+vehicle and MCAo+IESC secretome were examined for villus shape, such as blunting, broadening of the villus base, loss of the normal slender profile, and quantified as villus length-to-width ratio. In all cases, stroke caused a significant reduction of the villus length-to-width ratio as compared to sham, which was modified by treatment (IESC secretome), time after stroke (4d or 30d) or biological sex (male/female). In middle-aged females, MCAo produced a pronounced distortion of ileal morphology, (Fig. 1a, b), significantly reducing villus length-to-width ratio at both acute (Fig 1e) and chronic (Fig 1f) time points compared with sham controls. Administration of adult IESCs-secretome (3 doses) did not significantly reverse this structural injury at either time point (Fig 1e, f).

**Figure. 1.**
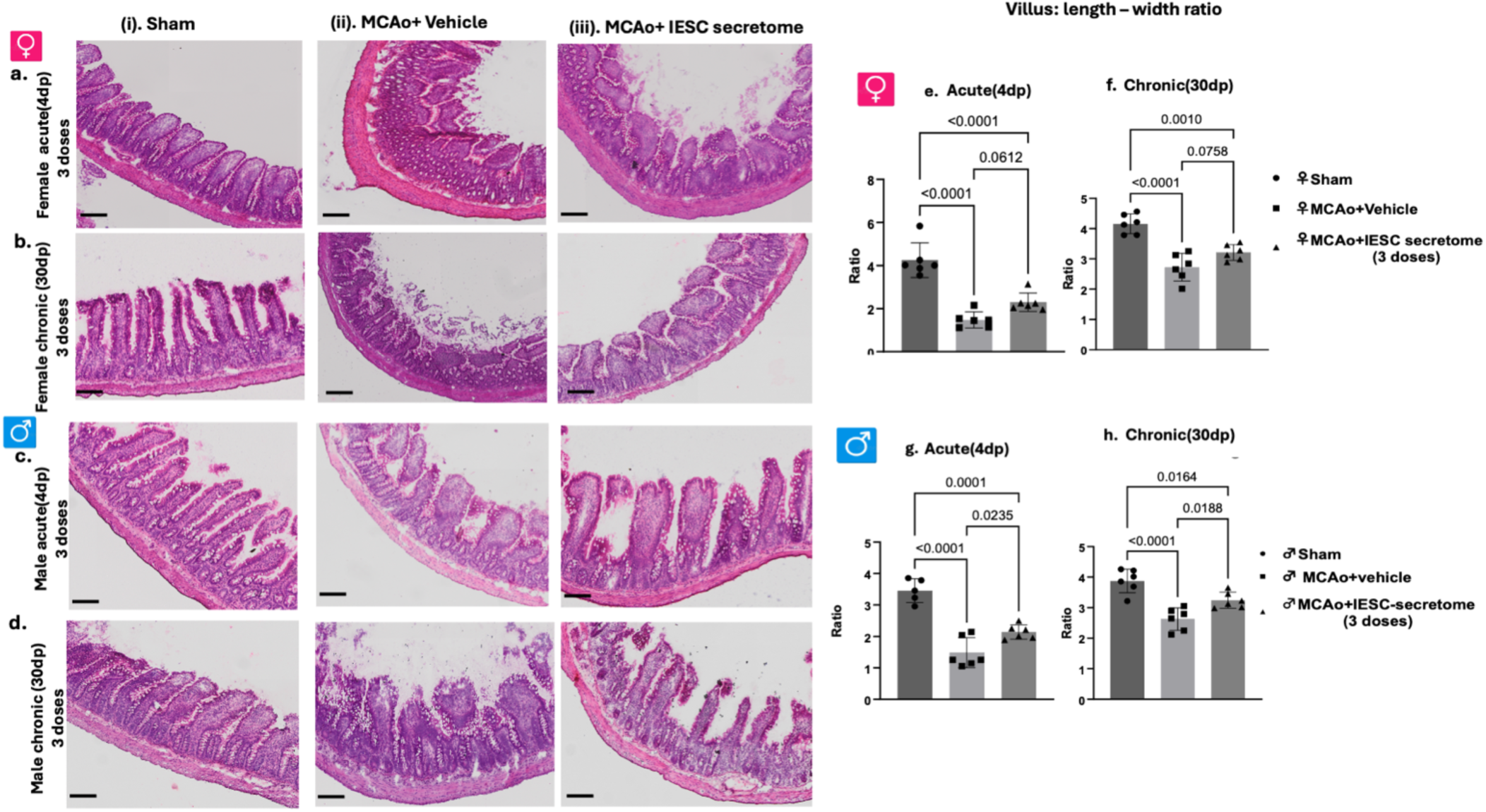
Effect of IESCs-secretome treatment on villus dysmorphia in the acute and chronic phase of stroke. Representative H&E-stained sections of small intestine (distal ileum) collected at 4 days and 30 days after MCAo from sham, MCAo+vehicle, and MCAo+IESC secretome treated females (a-b,(i-iii)) and males (c-d,(i-iii)). Histograms depict villus length-width ratio calculated from these sections (e-h). Data are presented as mean ± SEM; n = 5-6/group. Data were analyzed by one-way ANOVA with Tukey’s post hoc multiple comparisons test. Exact p values for pairwise comparisons are shown on the graph. Magnification bar: 32µm.

By contrast, middle-aged males exhibited clear structural improvement in response to IESC-secretome following MCAo. Vehicle-treated stroke males showed significant reduction in villus length-to-width ratio at both acute (Fig 1c, g) and chronic (Fig. 1d, h) time points compared with sham. Secretome-treated males displayed a robust increase in villus ratio relative to vehicle-treated MCAo animals in both the acute and chronic phase, although at both time points, stroke groups had lower ratios than shams, consistent with substantial treatment effect but incomplete long-term recovery of gut histoarchitecture.

### IESCs-secretome treatment differentially modulates stroke-induced goblet cell loss in the middle-aged rats

Gut epithelial structure was assessed by goblet cell counts, identified by PAS-positive staining, and reported as number of goblet cells per villus. Similar to villus morphology, goblet cell numbers (from ileum) were affected by stroke, treatment and biological sex. In middle-aged females, MCAo resulted in a marked reduction in goblet cell numbers per villus in both the acute (Fig. 2a) and chronic (Fig. 2b) phases compared with sham-operated controls. In the acute cohort, administration of three doses of adult IESCs-secretome did not significantly rescue goblet cell numbers relative to vehicle-treated stroke animals (Fig. 2e) and both stroke groups remained significantly below sham. In the chronic phase, secretome-treated female rats showed numerically higher goblet cell counts compared to vehicle-treated MCAo animals (Fig. 2f), but the groups were not statistically different and both stroke groups had lower goblet cell numbers relative to sham.

**Figure 2.**
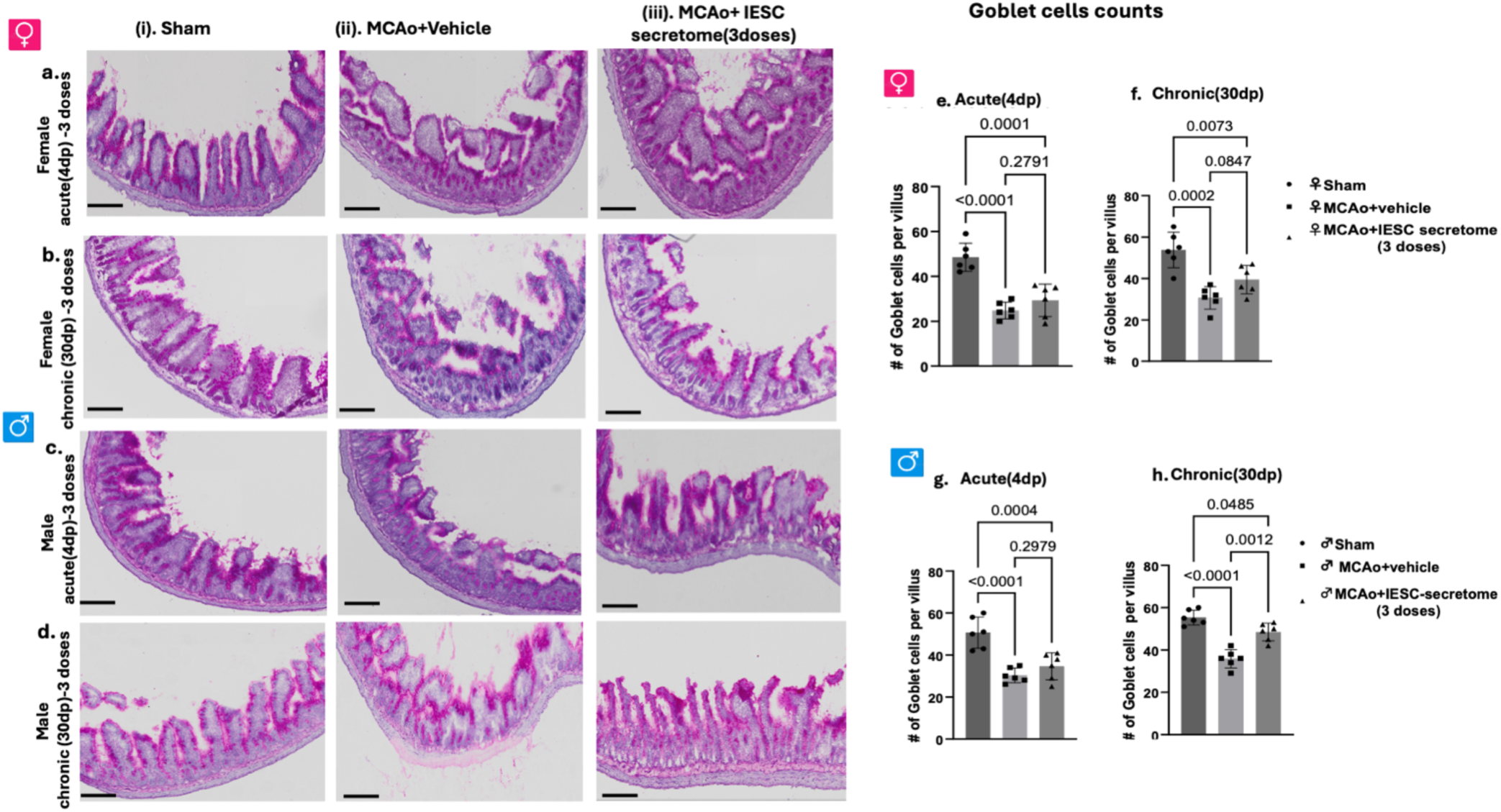
Effect of IESCs-secretome administration on goblet cell number in the acute and chronic phase of stroke. Representative images of Periodic acid–Schiff (PAS)-stained ileum sections obtained at 4 days and 30 days after MCAo from sham, MCAo + vehicle, and MCAo + IESCs-secretome treated females (a-b) and males (c-d). Histograms depict goblet cell number per villus calculated from PAS-stained sections (e-h). n= 5-6/group. Data are presented as mean ± SEM. Data were analyzed by one-way ANOVA with Tukey’s post hoc multiple comparisons test. Exact p values for pairwise comparisons are shown on the graph. Magnification bar: 32µm

In middle-aged males, MCAo similarly caused a robust decline in goblet cell numbers per villus at both acute (Fig. 2c) and chronic (Fig. 2d) time points compared with sham. In the acute phase, goblet cell counts were no different in secretome-treated animals and vehicle-treated MCAo rats (Fig. 2g). At the chronic time point, goblet cell numbers in secretome-treated males were significantly elevated relative to vehicle-treated stroke animals (Fig. 2h) and approached, but did not fully reach, sham levels. Overall, these findings indicate that a three-dose regimen of adult IESCs-secretome does not improve stroke-induced goblet cell loss in middle-aged females but promotes partial, delayed restoration of goblet cell density in middle aged males.

### IESC-derived secretome attenuates stroke-induced gut permeability

As a further measure of gut health, serum lipopolysaccharide (LPS) and intestinal fatty acid binding protein (iFABP), surrogate markers of endotoxemia and epithelial injury, respectively were assessed at 4 days and 30 days post-MCAo. LPS and iFABP levels were elevated by stroke at both time points. In middle-aged females, LPS levels were elevated in vehicle-treated MCAo female compared with sham controls (Fig. 3a; 27.2 ± 4.1 vs. 11.7 ± 1.8 ng/mL), while IESC-secretome administration significantly reduced circulating LPS levels to 16.1 ± 2.4 ng/mL. Male animals demonstrated a similar pattern, with vehicle-treated MCAo rats showing marked increases in LPS compared with sham (Fig. 3b; 59.3 ± 6.7 vs. 18.6 ± 2.3 ng/mL), which were significantly attenuated by IESC-secretome treatment (33.4 ± 4.2 ng/mL). Acute changes in iFABP mirrored those observed for LPS. In females, iFABP levels were significantly elevated in vehicle-treated MCAo animals (Fig. 3c; 161.7 ± 11.2 ng/mL) and were reduced by IESC-secretome administration to 104.2 ± 10.5 ng/mL. In males, MCAo induced a comparable increase in serum iFABP (Fig. 3d; 178.9 ± 9.6 ng/mL), which was significantly reduced following IESC-secretome treatment (126.3 ± 8.7 ng/mL).

**Figure 3.**
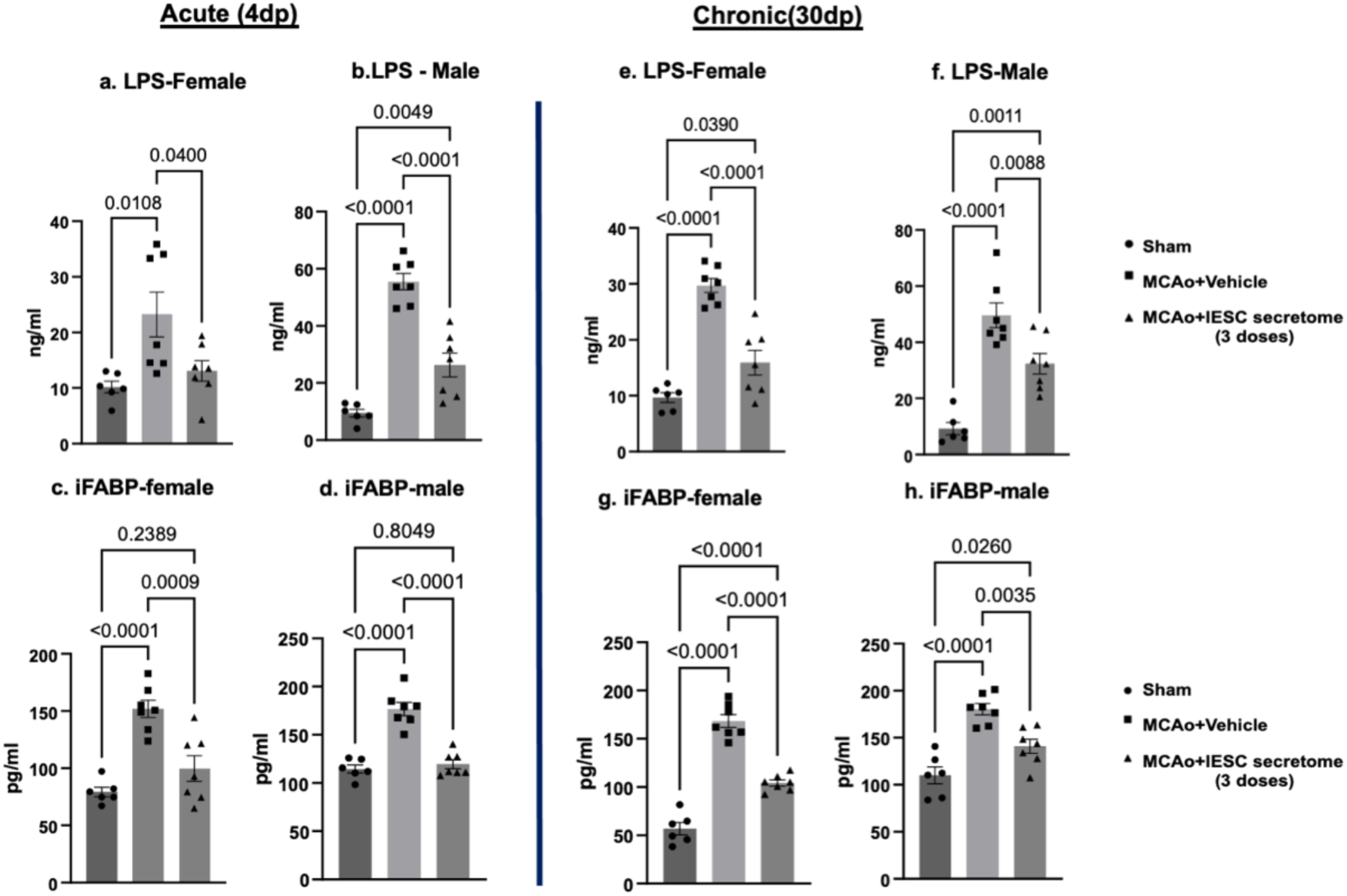
Effect of IESCs-secretome administration on serum LPS and iFABP as measures of gut permeability. Serum LPS and iFABP levels acute (4d post; left panel) and chronic (30d post; right panel) phase of stroke in females (a,c,e,g) and males (b,d,f,h). Histograms depict mean±SEM. N = 7-10/group. Data were analyzed by one-way ANOVA with Tukey’s multiple-comparisons test. Exact p values are shown on the graphs.

Persistent gut barrier dysfunction was evident 30d after MCAo in vehicle-treated animals of both sexes. In females LPS levels remained significantly elevated relative to sham (Fig. 3e; 32.6 ± 3.9 vs. 13.4 ± 2.2 ng/mL), while IESC-secretome treatment reduced LPS to near-sham levels (17.2 ± 2.7 ng/mL). In males, LPS levels remained elevated in the chronic phase and were similarly reduced by IESC-secretome administration (Fig. 3f; 61.1 ± 5.6 to 39.7 ± 4.1 ng/mL). Chronic iFABP levels followed the same trajectory, decreasing from 201.6 ± 12.3 to 127.3 ± 9.1 ng/mL in females (Fig. 3g) and from 188.5 ± 9.7 to 140.2 ± 7.6 ng/mL in males (Fig. 3h) following IESC-secretome treatment.

### IESC-secretome differentially modulates acute post-stroke peripheral cytokine responses in a sex-dependent manner

To assess whether adult IESC-derived secretomes alter stroke-induced systemic inflammation, circulating cytokines were quantified in serum from middle-aged male and female rats following MCAo. In females (Table 1), MCAo induced a robust pro-inflammatory response characterized by significant elevations in IL-17A, IL-6 and TNF-α relative to sham animals, however, IESC-secretome treatment did not attenuate these cytokines, and were no different from vehicle-treated animals. The anti-inflammatory cytokine IL-10 was markedly suppressed after stroke in both vehicle-and secretome-treated females, suggesting a persistent pro-inflammatory bias in the female stroke groups. In contrast, male animals exhibited a more treatment-responsive inflammatory profile. MCAo-induced elevation of circulating cytokines was significantly reduced by IESC-secretome treatment compared to vehicle treatment. IL-10 was suppressed after stroke as compared to sham although IESC-secretome treatment increased IL-10 levels. Together, these data demonstrate that adult IESC-derived secretomes exert sex-specific immunomodulatory effects following ischemic stroke, with greater anti-inflammatory efficacy in males.

**Table 1.** IESC secretome (3 doses) treatment on circulating cytokine levels at 4d after MCAo.

|  | Cytokine | Sham, mean ± SEM (pg/mL) | MCAo + Vehicle, mean ± SEM (pg/mL) | MCAo + IESC secretome, mean ± SEM (pg/mL) |
|---|---|---|---|---|
| **Female-acute(4dp)** | IL-17A | 68.4 ± 4.0 | 95.1 ± 4.6* | 86.9 ± 4.7 |
|  | IL-6 | 1089.1 ± 95.2 | 1713.3 ± 133.7* | 1453.6 ± 147.0 |
|  | TNF-α | 14.7 ± 2.4 | 27.4 ± 1.8* | 21.8 ± 1.6 |
|  | IL-10 | 419.2 ± 45.4 | 90.9 ± 8.5* | 127.9 ± 17.9 |
| **Male-acute (4dp)** | IL-17A | 101.3 ± 11.0 | 156.7 ± 6.5* | 118.5 ± 7.0** |
|  | IL-6 | 995.7 ± 57.3 | 2093.2 ± 98.2* | 1432.3 ± 138.2** |
|  | TNF-α | 24.0 ± 3.9 | 42.5 ± 2.0* | 28.5 ± 2.0** |
|  | IL-10 | 223.3 ± 12.5 | 117.6 ± 6.1* | 155.3 ± 7.5** |

Values shown are mean+SEM; \*: p<0.05 compared to Sham; \*\*:p<0.05 compared to MCAo+Vehicle

Overall, IESC secretome treatment after stroke improved histological features of the gut epithelium as well as reduced gut permeability and inflammatory cytokines in males, while in females, IESC secretome treatment improved gut permeability measures, but did not fully improve gut epithelium morphology or inflammatory cytokines.

### IESC-Secretome therapy improves acute stroke outcomes

We next evaluated the therapeutic potential of IESCs-secretome on acute stroke outcomes, including survival, infarct volume, sensory motor impairment (ART and VIB), and GFAP as a marker of blood brain barrier permeability. Survival: As shown in the Kaplan–Meier survival plots (Fig. 4a,g), MCAo-related mortality was reduced by IESC-secretome administration in both sexes. Among middle-aged female rats, IESC secretome treatment reduced mortality through 96 hours, although this was not statistically significantly different from vehicle treated animals (Fig. 4a). In middle-aged male MCAo rats, secretome treated also reduced mortality through 96 hours, and was significantly different from vehicle treated animals (Fig. 4g). Infarct volume, estimated from TTC-stained coronal brain sections obtained 4days post MCAo (Fig. 4b, h), was significantly reduced in IESC-secretome treated animals compared to vehicle-treated controls in both females and males. In females, the mean ratio decreased from 0.465 in the vehicle group to 0.210 in the IESC secretome group (Fig. 4c). Similarly, male animals showed a reduction from 0.544 to 0.162 following IESCs-secretome treatment (Fig. 4i). Serum glial fibrillary acidic protein (GFAP) levels were measured as an indicator of astroglial injury and BBB disruption. In females, MCAo induced a marked increase in GFAP levels in vehicle-treated animals as compared to sham, which was significantly reduced by IESC-secretome administration (Fig. 4d). A similar pattern was observed in males, where GFAP increased following MCAo and was significantly attenuated by IESC-secretome treatment (Fig. 4j).

**Figure 4.**
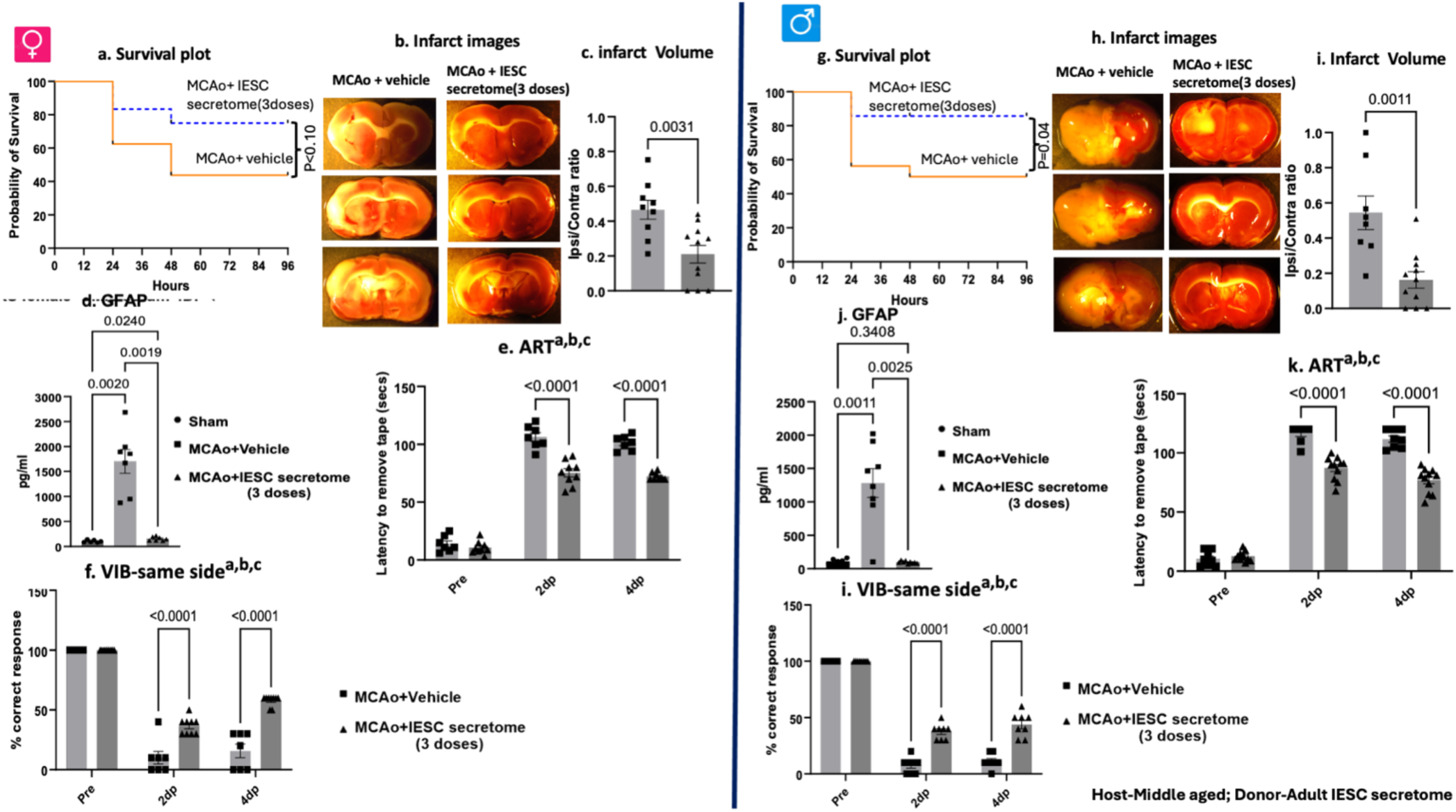
The effect of IESC-secretome treatment on acute stroke outcomes. Secretome from adult donors were administered to middle aged hosts at 4, 24, and 48h post MCAo. Female cohort shown on the left side panel (a-f), male cohort shown on the right side panel (g-l). a,g: Kaplan Meier survival plot depicting mortality after MCAo in vehicle-treated rats and IESC secretome treated females (a) and males (g). Representative TTC-stained coronal sections and quantitation of infarct volume from TTC-stained sections from vehicle and IESC-secretome treated females (b,c) and males (h,i). Serum GFAP levels were measured by ELISA in sham, MCAo + vehicle, and MCAo + IESCs-secretome treated females (d) and males (j). Sensory motor function was assessed pre stroke, 2d and 4d after stroke by the Adhesive Removal test (ART) and the same side VIB test in females (e,f) and males (k,i). Data were analyzed by ttests or two-way ANOVA coded for repeated measures followed by Tukey’s multiple-comparisons test. Key ^a^: main effect of stroke, ^b^: main effect of treatment, ^c^: interaction effect. n = 7-10/group. Exact p values for planned comparison as indicated on the graphs.

The acute neuroprotection was also reflected in better performance on sensory motor behaviors. In the adhesive tape removal test, MCAo resulted in longer latencies for tape removal at 2d and 4d post stroke (Fig. 4e, k), however IESC-secretome treated animals had significantly shorter latencies at both time points as compared to vehicle treatment. A similar effect of stroke and treatment was seen in the ‘same side’ vibrissae evoked forelimb placement task (Fig. 4f, i). Both vehicle-treated and secretome treated rats showed a significant deficit in paw placement reflex of the contralesional paw at 2 days and 4 days post stroke, however, IESC-secretome treated rats were less impaired than vehicle treated animals at both time points, indicating a beneficial effect of secretome.

Overall, this data provides strong evidence that IESC secretome delivery exerts neuroprotective effects in the acute post stroke period in both sexes. Despite the lack of immunomodulation of proinflammatory factors or effective gut epithelial repair in females, acute stroke outcomes were improved by IESC secretome treatment in both sexes.

To assess whether the improved acute stroke outcomes would also improve long term neurobehavioral we next assessed affective and cognitive behaviors 4+ weeks (30dp) after stroke.

### Social interaction

Administration of IESC secretome preserved social interaction behavior following ischemic stroke in both female and male middle-aged rats. In female rats, there was a significant effect of both treatment (IESC secretome vs vehicle) and time (pre vs post stroke). Vehicle-treated females exhibited a marked decline in time spent interacting with the social stimulus (stranger rat) after stroke (Fig. 5a). In contrast, IESC secretome-treated females maintained robust sociability across testing sessions, with no significant pre-to post-stroke difference, and significantly higher sociability than that of vehicle-treated controls.

**Figure 5.**
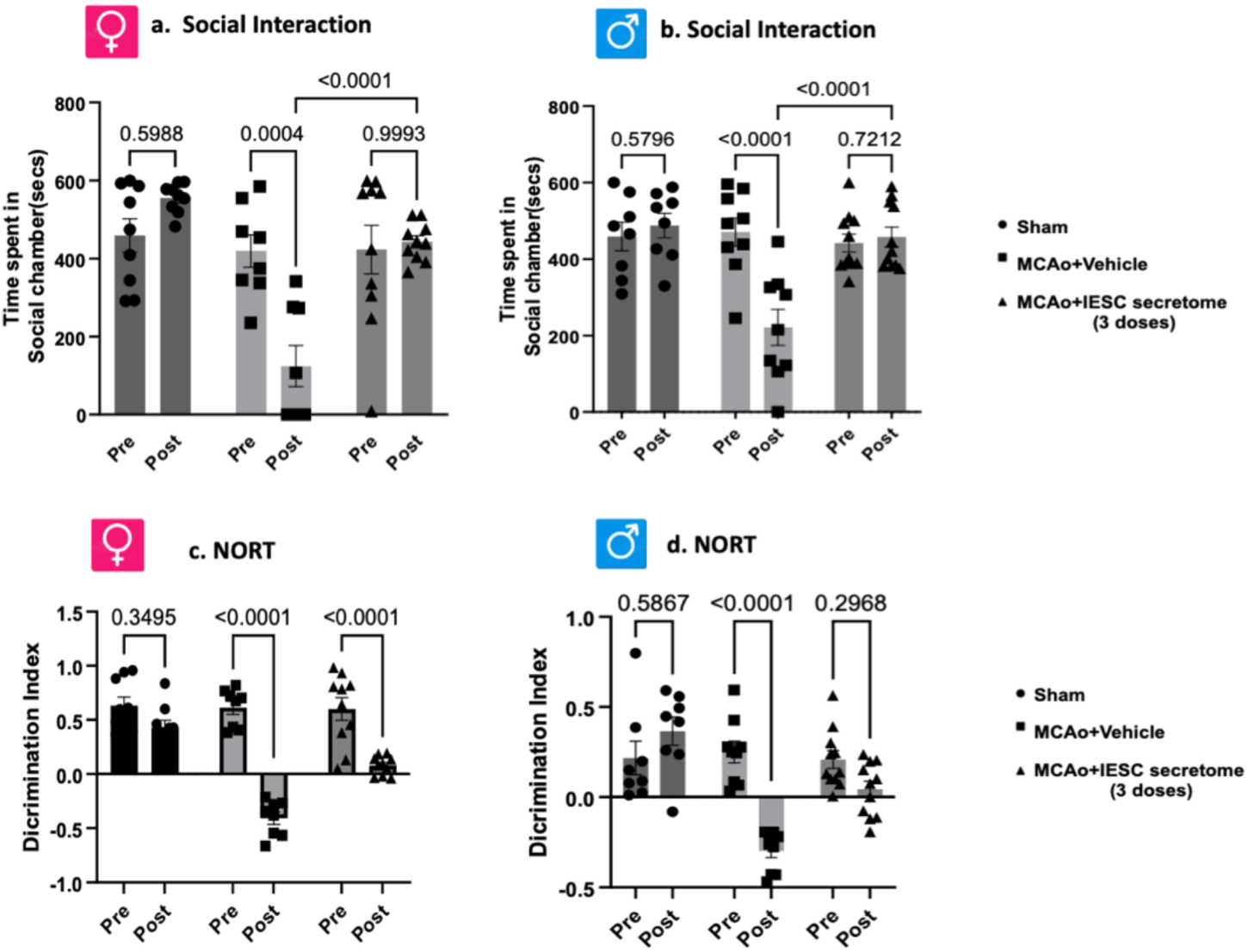
Effect of IESC secretome treatment on post-stroke social interaction behavior. (a, b) Time spent in the social interaction chamber was measured before (Pre) and after (Post) MCAo or sham surgery in middle-aged female (left) and male (right) rats treated with vehicle or adult IESC secretome. n = 7-10/group. Data are presented as mean ± SEM and analyzed by 2-way ANOVA with repeated measures followed by Tukey’s multiple-comparisons test. Exact p values for pairwise comparisons are shown on the graph. (c, d) Novel object recognition test (NORT) was used to assess episodic memory in females (left) and males (right) 30 days after stroke. Discrimination index (DI) was measured prior to stroke (Pre) and during the post-stroke testing phase (Post). The line at DI 0.0 on the Y axis indicates chance performance. n = 7-10/group. Data are presented as mean ± SEM and analyzed using 2-way ANOVA with repeated measures followed by Tukey’s multiple-comparisons test. Exact p values of pre and post comparisons are shown on the graph.

A similar protective pattern was observed in males. Stroke significantly reduced sociability in vehicle-treated male rats (Fig. 5b). However, males treated with IESC secretome displayed stable social interaction following stroke, and post-stroke sociability scores were markedly higher than those of their vehicle counterparts. Sham operated animals, regardless of sex, exhibited no significant changes in social interaction across time points, confirming that the observed effects were specific to ischemic injury and treatment intervention. These SI data reflect the emergence of stroke-induced social withdrawal and behavioral resilience in groups treated with adult IESC secretome.

### Post stroke cognitive impairment

#### Episodic memory

Episodic memory was assessed by the novel object recognition test (NORT) at 30 days post MCAo. Cognitive performance was quantified using the discrimination index (DI), where positive values denote a preference for the novel object. In vehicle-treated stroke females, the DI was significantly different 30 days after MCAo (Fig. 5c), with animals showing a preference for the familiar object. IESC secretome-treated females also differed significantly from their post stroke performance (Fig. 5c), although their discrimination index score was close to zero, indicating no preference for either novel or familiar object. Thus, IESC treatment did not preserve episodic memory in females. Similarly, vehicle-treated males exhibited a significant reduction in DI following stroke (Fig. 5d) and also showed a preference for the familiar object. IESC secretome-treated males did not show a significant difference in DI after stroke relative to pre-stroke. Sham operated animals of both sexes exhibited stable DIs across testing intervals, confirming that behavioral changes were stroke and treatment dependent.

### Spatial learning and retrieval

Spatial learning and retrieval (recall) was evaluated by the Barnes Maze task. This assay measures the animal’s ability to learn and recall the location of a hidden escape chamber using spatial cues. Learning, or the acquisition phase, was assessed over four consecutive training days, with escape latency (time to locate the escape box) serving as the primary measure of performance.

During the acquisition phase (learning), sham operated animals demonstrated effective learning, as indicated by significantly reduced escape latencies between Day 1 and Day 4 (Fig. 6a). In contrast, stroke impaired learning in vehicle-treated and IESC-secretome treated females, reflected by persistently high escape latencies across the training days. In males, sham operated animals showed effective learning with reduced escaped latencies between Day 1 and Day 4 (Fig. 6b). Vehicle-treated male rats failed to show improved latencies confirming the adverse effects of stroke on learning. In contrast, IESC secretome treatment male rats showed effective spatial learning, with shorter escape latencies over the training period (Fig.6b). Overall, sham and vehicle-treated males exhibited performance patterns similar to those observed in females, while secretome treatment was selectively beneficial in males but not females.

**Figure. 6.**
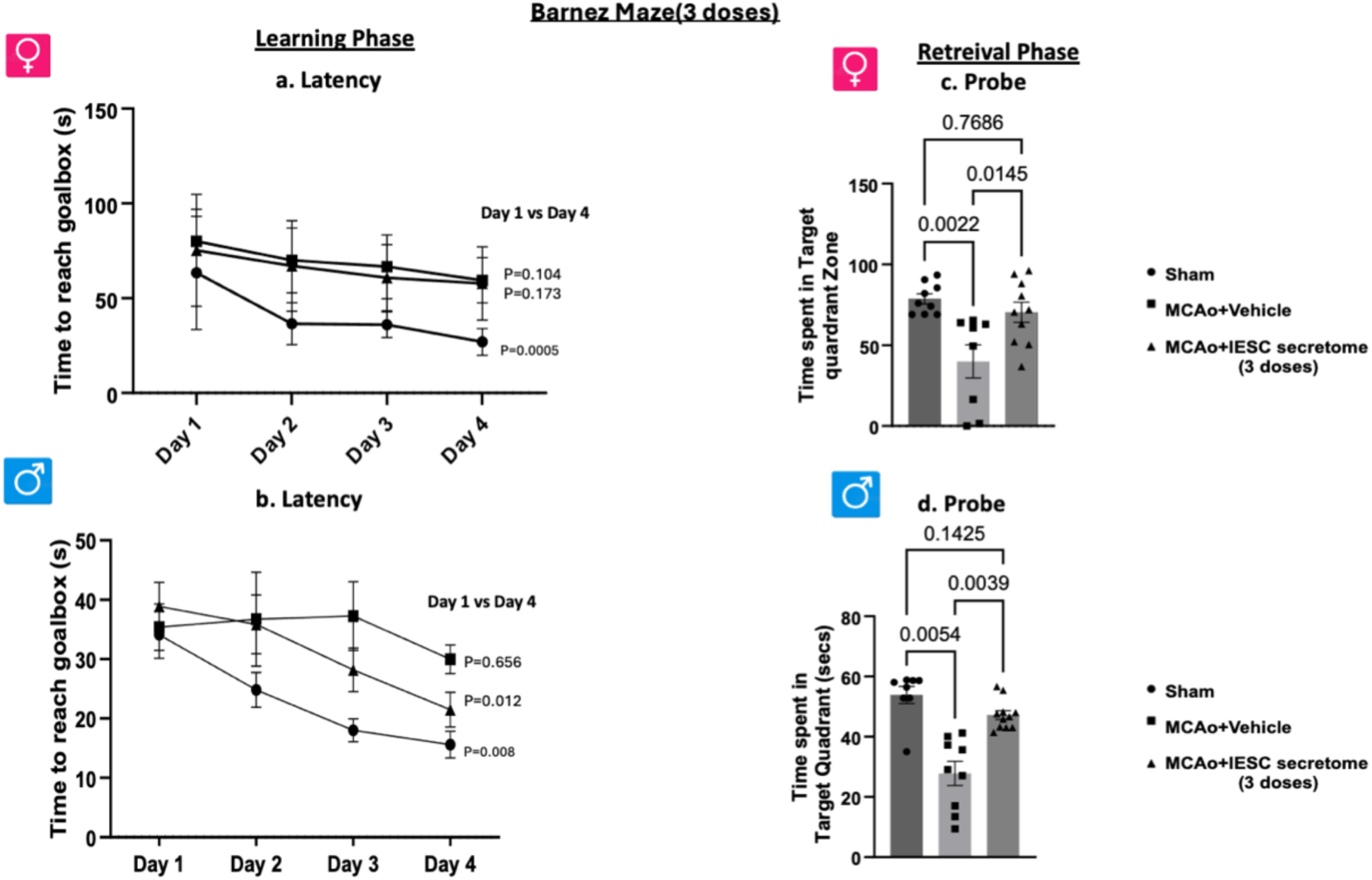
Effect IESC secretome treatment on spatial learning and recall after stroke. Spatial learning was assessed by latency to locate the escape hole across a 4-day acquisition period in Sham, MCAo+vehicle and MCAo+IESC secretome treated females (a) and males (b) 30 days after stroke. Latency on training Day 1 was compared with training Day 4 to assess effective spatial learning. Spatial reference memory was assessed by the probe trial 48 h after the last training day, as time spent in the target quadrant. n = 7-10/group. Data are presented as mean ± SEM. Group differences were assessed by one way ANOVA and Tukey’s multiple-comparisons test. For the learning phase, latency on day 1 was compared to latency on day 4. Exact p values shown on graphs.

Spatial retrieval was assessed by the probe trial, conducted 48 hours after the final training session. Longer times spent in the target quadrant is indicative of better recall. Vehicle-treated stroke animals, both male and female, spent significantly less time in the target quadrant than shams (Fig. 6c, 6d), while IESC treated males and females spent longer times in the target quadrant compared to vehicle treated animals, indicating preserved spatial recall. Importantly, both secretome-treated groups were not significantly different from their sham counterparts. Thus, while IESC secretome treatment was effective for preserving learning and recall in male stroke rats, secretome treatment was only effective for recall in female stroke rats.

Together, these results demonstrate that secretome treatment confers sex specific cognitive benefits following ischemic stroke and is consistent with the evidence that IESC treatment was less effective for gut repair (Fig. 1-3) and immune suppression (Table 1) in females as compared to males.

### Adult IESC-secretome treatment differentially modulates serum cytokines during the chronic phase of stroke

At the chronic time point, adult IESC-secretome produced modest, sex-dependent effects on circulating cytokines in middle-aged rats subjected to MCAo (Table 2). In females, MCAo+vehicle significantly increased IL-17A, IL-6, and TNF-α compared to sham-operated animals. IESC-secretome treated females did not differ from vehicle-treated females for any of these analytes. The anti-inflammatory cytokine IL-10 was decreased in MCAo-vehicle treated females compared with sham (p < 0.0001), as well as the secretome-treated group. These findings indicate that the three-dose adult IESC-secretome treatment exerts minimal corrective effects on the systemic cytokine profile in females, with persistent elevation of pro-inflammatory cytokines and suppression of IL-10.

**Table 2.** IESC secretome (3 doses) treatment on circulating cytokine levels at 30d after MCAo.

|  | Cytokine | Sham, mean ± SEM (pg/mL) | MCAo + Vehicle, mean ± SEM (pg/mL) | MCAo + IESC secretome, mean ± SEM (pg/mL) |
|---|---|---|---|---|
| **Female-Chronic(30dp)** | IL-17A | 105.3 ± 12.6 | 272.4 ± 37.6* | 253.5 ± 25.0 |
|  | IL-6 | 517.8 ± 91.6 | 1250.7 ± 152.6* | 1463.7 ± 125.7 |
|  | TNF-α | 18.1 ± 2.1 | 46.0 ± 3.0* | 37.8 ± 2.3 |
|  | IL-10 | 347.9 ± 25.3 | 121.9 ± 251* | 165.8 ± 20.6 |
| **Male-Chronic(30dp)** | IL-17A | 69.0 ± 4.3 | 142.2 ± 7.2* | 107.4 ± 7.6** |
|  | IL-6 | 697.5 ± 52.5 | 1637.4 ± 86.2* | 1015.2 ± 65.3** |
|  | TNF-α | 17.0 ± 1.8 | 31.3 ± 1.4* | 24.0 ± 2.0** |
|  | IL-10 | 423.2 ± 51.1 | 132.9 ± 11.9* | 270.3 ± 36.1** |

Values shown are mean+SEM; \*: p<0.05 compared to Sham; \*\*:p<0.05 compared to MCAo+Vehicle

In males (Table 2), chronic cytokine responses were more responsive to secretome therapy. Relative to sham animals, MCAo+vehicle significantly elevated IL-17A, IL-6 and TNF-α. In all cases, IESC-secretome treatment significantly reduced the levels of these cytokines compared to vehicle treatment, although they remained elevated compared to shams. In contrast, IL-10 was decreased by stroke in vehicle-treated males, while secretome treatment significantly increased this cytokine compared to vehicle, though both stroke groups were still lower than sham. Overall, these chronic-phase data demonstrate that adult IESC-secretome after stroke attenuates pro-inflammatory cytokines and enhances IL-10 in males, whereas females show little cytokine modulation in response to secretome treatment, reinforcing a sustained, sex-dependent difference in peripheral immune responsiveness to IESC-derived secreted factors.

### IESCs-secretome limits ischemic injury and reduces T2 hyperintensity volume after stroke in middle-aged rats

Postmortem T2-weighted MRI scans were obtained from middle-aged male and female stroke rats treated with either vehicle or sex-matched adult IESCs-secretome (Fig. S2a, c) The ratio of hyperintensity volume normalized to the non-ischemic hemisphere volume was markedly reduced by IESCs-secretome treatment compared with vehicle in females (Fig. S2b and males (Fig. S2d) further indicating substantial protection from ischemic edema and tissue swelling. This finding demonstrating that IESC secretome treatment reduced edema formation and tissue swelling in the chronic phase, although cognitive performance was impaired in females.

### Proteomic analysis of IESC secretome

Proteomic analysis of IESC secretome was prompted by the goal of identifying pathways that may confer stroke protection, with the additional goal of identifying sex differences in these pathways, based on the sex differences noted in the effects of IESC on gut repair and cognitive recovery.

We filtered canonical pathway results to include only those with an absolute activation z-score ≥ 2 and a Fisher’s exact test p-value < 0.05. These combined thresholds enrich pathways that are both significantly overrepresented in the IESC-secretome datasets and have a robust predicted direction of regulation. Shown in Fig. 7 are the top canonical pathways in males (A,B) and females (C,D). Each pathway is plotted as a bubble, the size of which indicates the number of genes in that pathway. In males, 19 of the 133 pathways met the criteria of a z-score ≥ 2. Predicted activation pathways, indicated by red bubbles, related to metabolic reprogramming (eg., glycolysis 1, gluconeogenesis1, glucose metabolism, peroxisomal lipid metabolism) signaling and cell regulation (eg., granzyme, GAIT translation, ephrin A signaling). In females, 20 of the 138 pathways met the criteria and predicted activation pathways included of protein synthesis (eg., Eukaryotic Translation Initiation, Eukaryotic Translation Elongation, Eukaryotic Translation Termination, Ribosomal Quality Control Signaling Pathway Response of EIF2AK4 (GCN2) to amino acid deficiency), protein modification and degradation (eg., Protein Ubiquitination Pathway, Deubiquitination, Neddylation), gene expression and RNA processing (Processing of Capped Intron-Containing Pre-mRNA, Major pathway of rRNA processing in the nucleolus and cytosol, MicroRNA Biogenesis Signaling Pathway, Nonsense-Mediated Decay (NMD)), signaling pathways (Signaling by ROBO receptors, Exosome Signaling Pathway, RAF/MAP kinase cascade, Signaling by NOTCH4).

**Figure 7.**
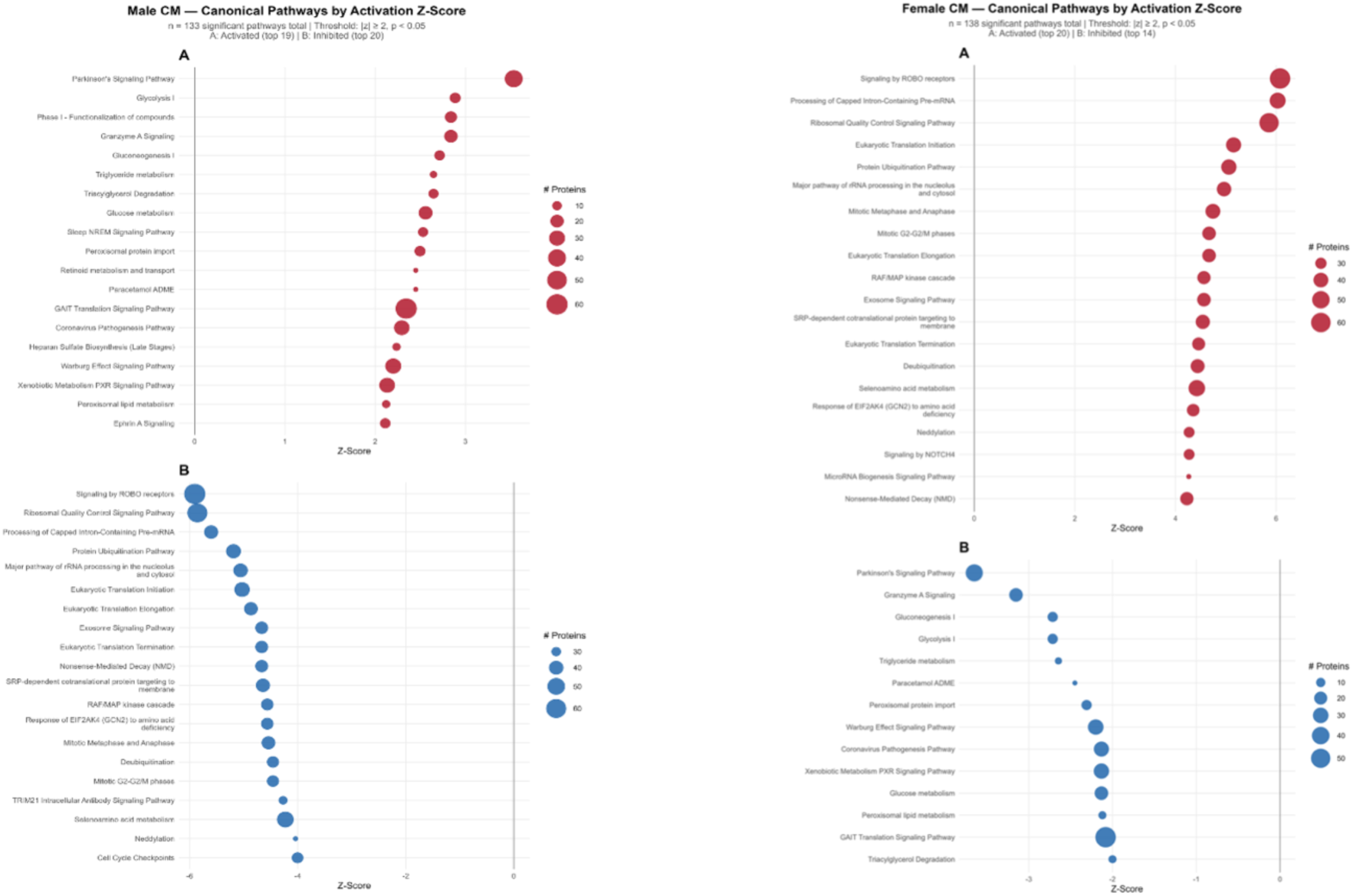
Analysis of Top Canonical pathways in male and female secretome. A, B: Canonical pathways significantly associated with the male-derived secretome (Fisher’s exact p < 0.05, |activation z-score| ≥ 2), out of 133 total significant pathways identified. (A) 19 pathways predicted to be activated (z-score ≥ 2). (B) Top 20 pathways predicted to be inhibited (z-score < −2). C, D: Canonical pathways significantly associated with the female-derived secretome (Fisher’s exact p < 0.05, |activation z-score| ≥ 2), out of 138 total significant pathways identified. (C) Top 20 pathways predicted to be activated (z-score ≥ 2). (D) 14 pathways predicted to be inhibited (z-score < −2). Dot size represents the number of proteins from the submitted dataset mapping to each pathway. Pathway analysis was performed using Ingenuity Pathway Analysis (IPA; QIAGEN) with the biological filter set to Rattus norvegicus.

Overall, sex differences in the IESC secretome arise primarily from differences in pathway-level regulation, rather than from wholesale gain or loss of specific pathways.

### Sex-dependent differences in selected IESC derived secretome growth factors

In addition to those detected by proteomic analysis, the expression of specific low molecular weight growth factors were assayed in the IESC-derived secretome. ELISAs were performed to measure IGF-1 VEGF-A, BDNF and PDGFa in adult female-and male-derived IESC secretome. While all were detected in male and female secretome, IGF-1 (Fig. S3a) and VEGF (Fig. S3b) concentrations were higher in females, while BDNF (Fig. S3c) was higher in males. PDGF-a (Fig. S3d) concentrations did not differ significantly between the sexes. Thus PDGF-a represents a conserved component of the IESC secretome, while IGF-1, VEGF-A, and BDNF show sex specific enrichment, suggesting the potential for differential modulation of angiogenic and neurotrophic signaling pathways.

### 4-doses IESC-secretome: effect on intestinal architecture and goblet cell density

Reconciling the broad commonalities in the male and female secretome pathways, with the sex differences in the effect of each secretome on gut and inflammatory responses, and long term behavioral outcomes, we next investigated whether increasing the number of doses of the female secretome (from 3 doses to 4 doses) would improve functional recovery in middle-aged female hosts. Both acute and chronic time points were evaluated in the 4 doses regimen which spanned 4h/12h/24h/36h after MCAo. A critical goal was to first determine whether the 4-dose regimen improved intestinal histoarchitecture, restored goblet cell populations and attenuated inflammatory cytokines compared to the earlier, less efficacious, 3-doses paradigm in females.

At the acute time point, vehicle-treated MCAo females showed pronounced villus blunting, epithelial disorganization (Fig. 8a) and goblet cell depletion (Fig. 8c) compared with sham controls while the 4-dose IESC-secretome administration visibly restored villus architecture and crypt-villus organization in stroke animals. Quantitative analysis of villus length:width ratio was significantly reduced in vehicle-treated animals compared with sham (Fig. 8e), whereas secretome treatment significantly increased this ratio compared to vehicle-treated animals. Goblet cell counts per villus were also markedly decreased in acute vehicle-treated MCAo rats relative to sham (Fig. 8g) but were rescued by 4 dose IESC-secretome. In both cases, secretome treated animals were significantly different from sham (Fig. 8e, g), indicating incomplete restoration of mucin-producing cells in the acute phase.

**Figure 8.**
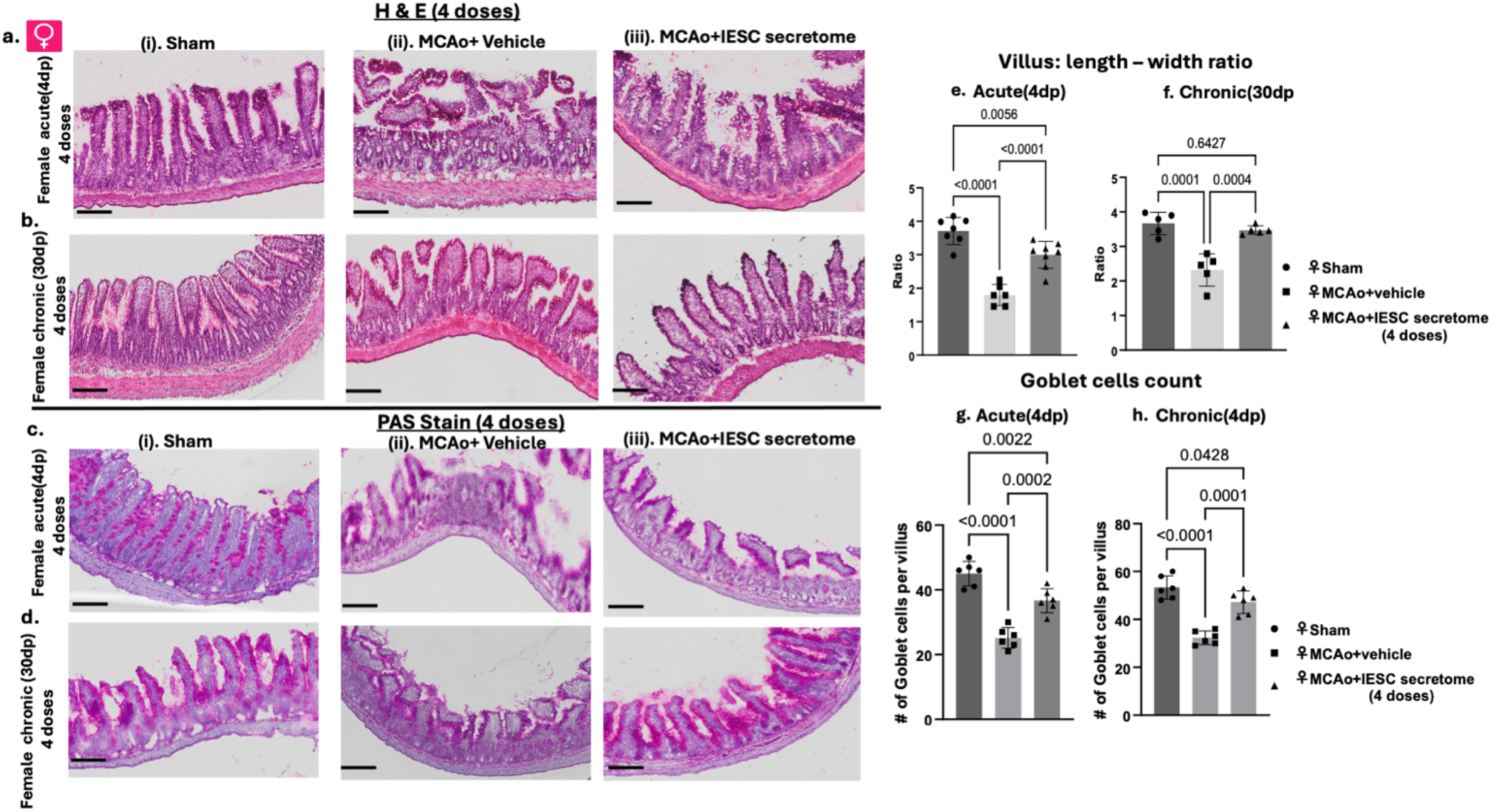
Effect of enhanced (4 doses) IESC secretome treatment on stroke-related gut dysmorphia and goblet cells. Representative H&E-stained sections of the ileum (a,b) and PAS-stained sections (c,d) from sham, MCAo + vehicle, and MCAo + IESCs-secretome (4 doses)-treated middle-aged female rats. Histogram of villus length-width ratio was calculated from H&E-stained images at the acute (e) and chronic (f) time points. Histogram of the number of goblet cells per villus were quantified from PAS-stained images at the acute (g) and chronic (h) phases. n = 5-6/group. Data are presented as mean ± SEM. Group differences were determined by one-way ANOVA with Tukey’s post hoc multiple comparisons test. Exact p values are shown on the graphs.

Persistent gut pathology was evident at the chronic time point (30dp) in vehicle-treated stroke animals compared with sham intestines (Fig. 8b). In contrast, animals receiving 4-dose IESC-secretome demonstrated improvement in mucosal morphology, with elongated villi and more organized epithelial layering that closely resembled sham sections. Quantitatively, villus length: width ratio remained significantly lower in chronic vehicle-treated animals as compared to sham (Fig. 8f), whereas secretome-treated rats exhibited a higher ratio than vehicle treated animals and were statistically similar to sham, indicating near-complete normalization of villus shape. Goblet cell numbers in the chronic phase followed a similar pattern; counts per villus were significantly reduced in vehicle-treated animals compared with sham (Fig. 8d) but were significantly increased by IESC-secretome treatment and statistically indistinguishable from sham (Fig. 8h), demonstrating restoration of mucosal secretory capacity.

To determine if the 4-dose IESC-secretome shapes the initial systemic inflammatory response to stroke, serum cytokines were assessed at 4 days and 30 days post-ischemia. In the acute cohort, MCAo +vehicle treatment (Table 3) produced a robust pro-inflammatory shift relative to sham controls, with marked increases in circulating IL-17A, IL-6, and TNF-α. The 4-dose secretome treatment significantly attenuated this surge in inflammatory cytokines compared with vehicle-treated MCAo animals. In parallel, the anti-inflammatory cytokine IL-10 was significantly decreased in vehicle-treated stroke animals relative to sham, while the 4-dose secretome administration elevated IL-10 concentrations.

**Table 3.** IESC secretome (4 doses) treatment on circulating cytokine levels.

| **Table-3** | **Cytokine** | **Sham, mean ± SEM (pg/mL)** | **MCAo + Vehicle, mean ± SEM (pg/mL)** | **MCAo + IESC secretome, mean ± SEM (pg/mL)** |
|---|---|---|---|---|
| **Acute(4dp)-female** | IL-17A | 54.3 ± 5.4 | 218.9 ± 20.3* | 91.7 ± 5.6** |
|  | IL-6 | 764.9 ± 171.0 | 1991.8 ± 326.3* | 902.9 ± 107.7** |
|  | TNF-α | 4.7 ± 0.9 | 23.6 ± 1.7* | 10.0 ± 0.7** |
|  | IL-10 | 278.8 ± 24.6 | 115.3 ± 25.4* | 198.5 ± 8.1** |
| **Chronic(30dp)-Female** | IL-17A | 51.5 ± 2.3 | 143.6 ± 9.7* | 74.0 ± 5.6** |
|  | IL-6 | 652.4 ± 35.8 | 1923.1 ± 150.4* | 940.5 ± 26.0** |
|  | TNF-α | 7.8 ± 0.6 | 21.4 ± 1.5* | 11.6 ± 0.7** |
|  | IL-10 | 257.5 ± 24.6 | 81.2 ± 5.3* | 140.6 ± 9.9** |

Values shown are mean+SEM; \*: p<0.05 compared to Sham; \*\*:p<0.05 compared to MCAo+Vehicle

At 30 days post-MCAo, vehicle-treated stroke animals exhibited a persistent pro-inflammatory milieu compared with sham controls, and marked reductions in IL-10. The 4-dose IESC-secretome administration robustly attenuated this chronic inflammatory profile relative to vehicle, while significantly increasing IL-10. Together, these data indicate that early, repeated IESC-secretome treatment reprograms long-term systemic immune signaling after stroke, producing durable attenuation of pro-inflammatory cytokines and partial restoration of anti-inflammatory mediators.

Overall, the 4-dose IESC-secretome treatment stabilized intestinal structure and barrier-supporting goblet cell populations and substantial dampening of systemic inflammation in middle-aged females, suggesting that this regimen may improve acute and chronic stroke endpoints.

### Effect of 4-dose IESC secretome treatment on acute stroke outcomes

Acute stroke: The 4 dose secretome treatment reduced stroke-related mortality as compared to MCAo+vehicle treated animals (Fig. 9a, Kaplan–Meier analysis, p=0.0415) restoring survival rates to those observed in sham-operated animals. Infarct volume (Fig. 9b) was significantly decreased in the 4-dose IESC group compared to the vehicle treated group (Fig. 9c). Latency in the adhesive removal test (ART), which was significantly impaired in the MCAo+vehicle treated animals at both 2 and 4 days (Fig. 9d) was attenuated by IESCs secretome treatment. Similarly, in the vibrissae-evoked forelimb placing (VIB) task, MCAo + vehicle animals exhibited profound deficits in same-side forelimb placing at 2 and 4 days post-stroke (Fig. 9e), while this deficit was attenuated in the 4-dose IESC-secretome group.

**Figure 9.**
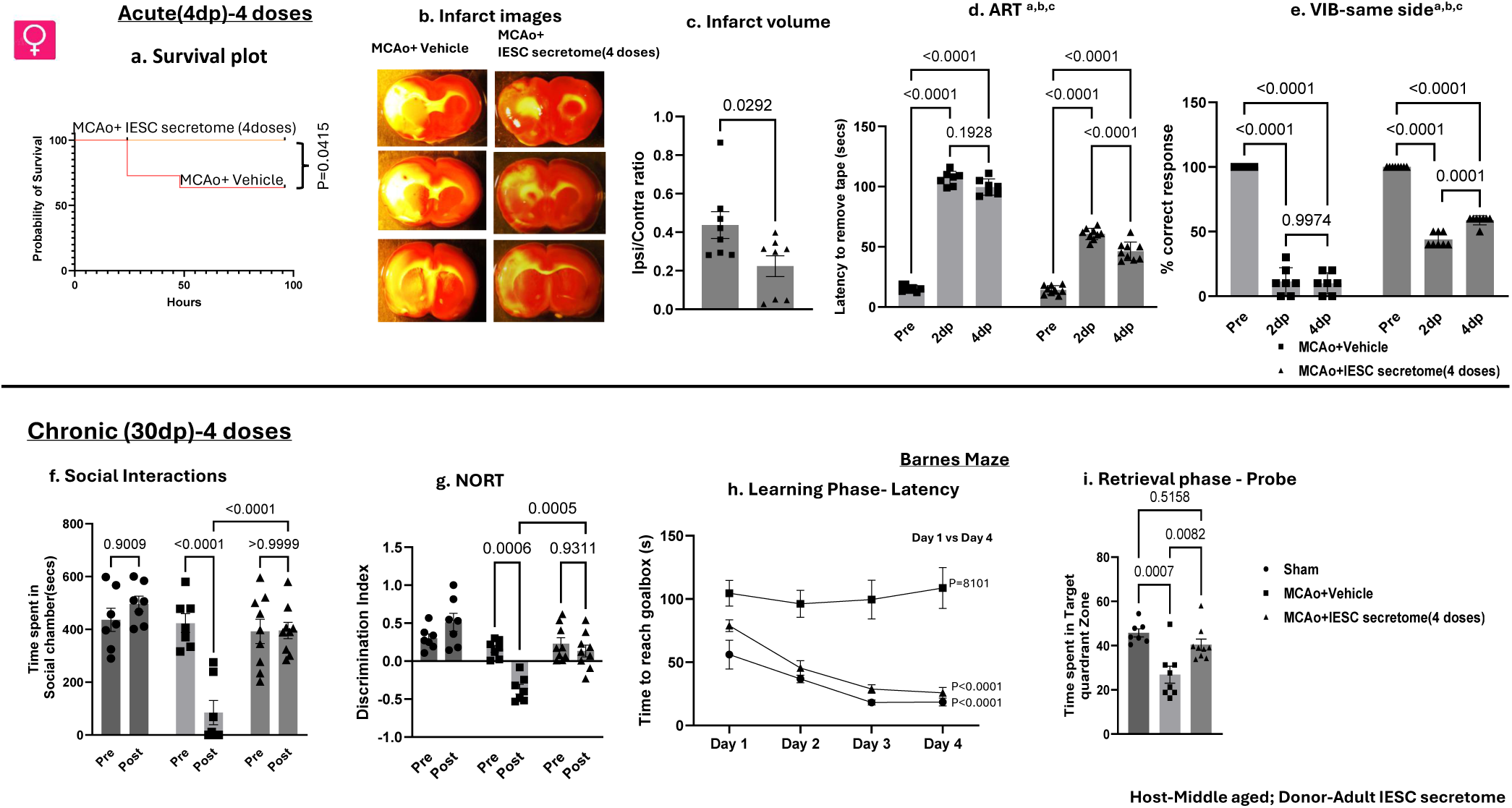
The effect of enhanced (4 doses) IESC-secretome treatment on acute and chronic stroke outcomes. **Acute study (a-e):** Kaplan Meier survival plot depicting mortality after MCAo in vehicle-treated rats and IESC secretome treated females. Representative TTC-stained coronal sections and quantitation of infarct volume from TTC-stained sections from vehicle and IESC-secretome treated females (b,c). Sensory motor function was assessed pre stroke, 2d and 4d after stroke by the Adhesive Removal test (ART) (d) and the same side VIB test (e). Key ^a^: main effect of stroke, ^b^: main effect of treatment, ^c^: interaction effect. Statistical analyses were performed using Two-way ANOVA followed by Tukey multiple-comparisons test. Graphs depict mean ± SEM. n= 8/group. Exact p values are shown on the graphs. **Chronic study (f-i)**: (a). Social interaction: Time spent in the social interaction chamber (Pre) and after (Post) MCAo or sham surgery in middle-aged female rats treated with vehicle or IESC secretome. (b) Novel object recognition test (NORT) was used to assess episodic memory in sham, MCAo+vehicle and MCAo+IESC secretome treated female animals. Discrimination index (DI) was measured prior to stroke (Pre) and during the post-stroke testing phase (Post). The line at DI 0.0 on the Y axis indicates chance performance. (c) Spatial learning was assessed by latency to locate the escape hole across a 4-day acquisition period in the Barnes maze test. For each group, p value shown compares latency on day1 with day 4. (d) Spatial reference memory was assessed by the probe trial 48 h after the last training day on the Barnes maze. Graph depicts time spent in the target quadrant where the escape hole was previously located. Data are presented as mean ± SEM; n = 7-10/group. For Social interaction and NORT, data were analyzed using 2-way ANOVA with repeated measures followed by Tukey’s multiple-comparisons test. Barnes maze learning and recall was measured by one-way ANOVA measures followed by Tukey’s multiple-comparisons test. Exact values of pre and post comparisons are shown on the graph.

Collectively, the acute phase findings in the 4-doses study mirrors that of the 3-doses study, showing substantially reduced infarct burden, improved survival, and accelerated sensorimotor recovery in middle-aged female rats.

### Chronic phase (4 doses): affective and cognitive impairments

We next examined whether this early immunomodulatory intervention translated into long-term behavioral and cognitive benefits. *Social interaction*: Baseline (pre-stroke) social interaction times did not differ across Sham, MCAo + IESCs-secretome groups (Fig. 9f). Following stroke, vehicle-treated stroke females exhibited a profound reduction in time spent in the social chamber compared with their pre-stroke performance (Fig. 9f). In contrast, the 4-dose IESCs-secretome treated animals maintained social interaction times comparable to their pre-stroke times and comparable to sham animals.

#### Novel object recognition test (NORT)

At baseline, discrimination indices (DI) were comparable across sham and stroke groups (Fig. 9g). Post-stroke, vehicle-treated animals exhibited significantly reduced DI relative to their own pre-stroke performance. In the 4-dose IESCs-secretome treated group, post stroke DI was indistinguishable from pre-stroke values.

#### Barnes maze acquisition (Learning phase)

Vehicle-treated animals displayed no improvement in latency to find the escape box over the training days (Fig. 9h) indicative of impaired task engagement. In contrast, the 4-dose IESC-secretome treated animals exhibited progressive improvement in latency across training days that closely paralleled the performance of Sham animals.

#### Barnes maze retrieval phase (recall)

In the probe trial, vehicle-treated animals spent significantly less time in the target quadrant compared with Sham controls (Fig. 9i), while the 4 dose IESCs-secretome treated animals exhibited longer target quadrant occupancy relative to vehicle and did not differ significantly from Sham animals, demonstrating effective restoration of spatial memory following this treatment regimen.

### Principal Component Analysis (PCA): Gut Barrier, Inflammatory, and Behavioral Profiles

PCA, applied to selected acute and chronic outcomes, indicated a clear separation between Sham and Vehicle-treated stroke animals along PC1, with Secretome-treated animals occupying an intermediate position. In the acute dataset (Fig. S4a-c), this separation was driven by opposing loadings of gut barrier markers (villus length-to-width ratio, goblet cell count) and pro-inflammatory cytokines (IL-6, IL-17a, TNF-α) across all cohorts. In the chronic dataset (Fig. S4d-f), separation for vehicle animals were driven by higher latency in the Barnes maze test (final training day), and lower delta in social interaction times and NORT preferences. Across all acute biomarkers (Fig. S4a-c), secretome treatment shifted the biomarker profile toward the Sham (healthy) condition, with the magnitude of this effect consistently greatest in females receiving four doses, followed by males receiving three doses, and weakest in females receiving three doses. This was reflected in the chronic outcomes as well, consistent with our hypothesis that acute gut histopathology and inflammatory mediators associate with long term neurobehavioral outcomes.

## DISCUSSION

This study demonstrates that the secretome of intestinal epithelial stem cells (IESC) is sufficient to improve stroke outcomes in middle-aged rats. Sex-matched adult donor IESC secretome administered intraperitoneally to middle-aged hosts after stroke improved both acute and chronic stroke outcomes, indicating that gut-derived paracrine factors can protect the injured brain without requiring cell transplantation. Secretome-based neuroprotection is consistent with a broader literature showing that stem cell conditioned media and extracellular vesicles improve outcomes after cerebral ischemia. Systemic delivery of mesenchymal stem (stromal) cell exosomes or extracellular vesicles, for example, promote functional recovery, enhance neurovascular plasticity, and reduce white matter damage after stroke in rodents (22, 29, 30). The present data therefore place adult IESC secretome alongside these other acellular products as a potent modulator of post-stroke injury and recovery. The proteome of the adult donor IESC secretome, the majority of which were common to both sexes, indicated pathways enriched for carbon metabolism, TCA cycle, pyruvate and fatty acid metabolism, and mitochondrial translation pathways, indicating common metabolic and translational components, a pattern seen in other stem cell-derived secretomes (31, 32).

Despite the commonalities in the conserved core proteome of the male and female secretome, we observed unexpected sex differences in treatment efficacy in the host, especially in long-tern behaviors. Sex-based transfer of donor secretome to middle aged stroke hosts significantly reduced acute stroke outcomes, such as infarct volume and sensorimotor impairment in both female and male rats. However, in males, secretome-treatment resulted in a near-complete rescue of social interaction behavior as well as episodic and spatial memory. In females, secretome treatment rescued social interaction behavior, but did not improve novel object recognition or spatial learning. These data point to a potential shortcoming of preclinical assessments. Based on the assumption that reducing the loss of brain tissue in the early aftermath of ischemia would limit subsequent chronic impairment, most preclinical stroke studies typically use acute phase outcomes to assess the new stroke neuroprotectants. However recent work from our lab has shown that infarct volume is not proportionally related to long term cognitive impairment. For example, intracerebroventricular administration of the peptide growth factor IGF-1 resulted in robust neuroprotection in the acute phase but did not improve stroke-induced cognitive impairment in the chronic phase (19). Infarct volume and clinical outcomes in patients from the ESCAPE-NA1 study reported that in very large infarcts (>200 ml), infarct volume was the leading determinant of outcomes, while in small infarct volumes (0-15ml) there was virtually no relationship with clinical outcomes(33). The current study thus adds to the literature that acute infarct volume is not predictive of long-term outcomes.

The present study suggests that gut health and circulating inflammatory mediators may be more accurate predictor of long-term outcomes than acute phase infarct volume. Thus, in middle-aged females, while infarct volume was attenuated by the three dose secretome treatment, villus architecture and goblet cell density failed to normalize and systemic cytokine profiling showed elevated levels of pro-inflammatory cytokines (IL-17a, IL-6, and TNF-α) and suppression of IL-10. These findings suggest that acute neuroprotection and chronic phase cognitive improvement are likely mediated by different factors, with persistent systemic and intestinal pathology more likely to impact cognitive function. This hypothesis is consistent with data from males, where the secretome treatment effectively reduced systemic inflammatory mediators (cytokines, LPS) and repaired villus structure and ameliorated cognitive performance.

This unexpected sex difference in treatment efficacy, especially in the context of inflammatory cytokines levels, led us to reconsider the dosing strategy. The three dose strategy (4, 24 and 48h after MCAo) was adopted based on the effectiveness of our previous cell transplant study, where IESCs were administered at the same three timepoints. Subsequent studies using an enhanced four dose secretome regimen in female hosts produced a functionally different outcome. This regimen induced coordinated modification of circulating and peritoneal cytokines, including marked reductions in endotoxin, IL-6, IL-17a, and TNF-α and partial restoration of IL-10, with full restoration of villus architecture and goblet cell density at both acute and chronic time points. Long-term social interaction, recognition memory, and Barnes maze probe performance were normalized to sham-like levels, indicating durable protection across affective and cognitive domains. Together, these findings support our assertion that gut repair and modification of key cytokines is a prerequisite for improving long term stroke.

Two hypotheses may account for the effectiveness of enhanced secretome treatment in females. One possibility is that, compared to males, the female secretome is deficient in trophic proteins and the additional dose compensates for this low abundance. Proteomic analysis of the secretome indicated that although there were sex-specific variations, there is a largely conserved core protein ensemble. For instance, both male and female secretomes contain growth factors with well-documented role in stroke, such as BDNF, a neurotrophin which promotes neuronal survival and functional recovery (34, 35), IGF-1 and VEGF-A which play a role in angiogenesis, neuronal survival, and functional recovery after experimental stroke (36–39), albeit expressed at different levels in male and females secretomes. Overall, the data from the four doses treatment suggests that the female proteome is not lacking in trophic support. A second possibility is that the timing of the additional dose is the critical determinant, aligning trophic availability with evolving stroke pathology. Stroke pathophysiology includes central and peripheral immune activation, which contributes to the extent of brain injury as well as long term behavioral impairment. In most cases, this response is similar in young and aged males and females, however, sex differences have been reported in middle age. At 24h after reperfusion in mice, middle aged male mice had elevated levels of CD45+high/CD3+ T cells compared to age matched females, while dendritic cells and Gr1+ macrophages were elevated in the middle-aged female brain compared to age matched males (40). While there are multiple peripheral sources for these cells, it is worth noting that dendritic cells were elevated in the spleen of females compared to males. Thus, it is likely that the early bulk secretome dosing in middle aged females in this study may have dampened an early inflammatory response resulting in improved long-term recovery.

Gut histology and inflammatory cytokines appear to be the key determinants that associate IESC secretome treatment with the potential for improvement in cognitive-affective outcomes. In the case of inflammatory cytokines, considerable evidence shows that these mediators are linked to depression (41) and post stroke cognitive impairment (PSCI) (42) as well as neurodegenerative diseases such as Alzheimer’s Disease (43, Swardfager, 2010 #108, 44). For example, IL-6 is positively correlated with stroke volume and a less favorable 1 year prognosis (45) and is inversely correlated with MMSE scores (46). IL-17A has been implicated in multiple human and animal stroke studies where it has also been shown to be a mediator of tissue damage (47). The link to gut histology, acute neurological injury and inflammatory is well-studied in the case of IL-17A, the characteristic effector cytokine of Th17 and γδ T cells. TH17/ γδ T cells are located in mucosal barrier tissues including the lamina propria of the intestinal tract Stroke induced intestinal barrier breakdown drives Th17/γδ T-cell activation and extravsation to the (22, 42, 43) (48). Although we cannot definitively state that elevated circulating levels of IL-17A is due to extravasation of gut resident Th17 cells, treatments that restore the barrier concomitantly decrease circulating IL-17A, as seen in this study, as well as our previous work using IESC transplants (23) and the peptide hormone IGF-1 (24).

Taken together, this work demonstrates that the secretome of adult IESC is a potent, gut focused paracrine therapy for stroke that can reduce infarct size, preserve barrier integrity, and improve complex behaviors, but whose efficacy is tightly constrained by sex, dose, and timing. By stabilizing the intestinal epithelium, lowering endotoxemia, and reprogramming cytokine networks, secretome treatment transects a pathological stroke-gut-brain loop and, under an early and intensive dosing regimen, achieves functional benefits comparable to IESC cell transplantation. The need for higher dosing in middle-aged females, highlight donor sex as a significant parameter for translating this approach. These findings support the development of standardized, rejuvenated IESC-derived secretome products as off-the-shelf biologics that could be combined with existing reperfusion therapies and targeted immunomodulation to extend the therapeutic window and improve long-term recovery after stroke.

## DECLARATIONS

### Author Contributions

KKM co-conceived the study, acquired funding support, coordinated all experiments, data acquisition and analysis, manuscript drafting and revision. AAM performed the proteomics pathway analysis. KP, AO, KAM ASK, NA, SRJ contributed to tissue collection, tissue processing for sectioning and staining, and assisted in behavioral assays and data analysis in 3 doses study; ZA contributed to perfusion, sample collection and imaging in the 4 doses study; ES, NA, NO, SM contributed to the animal behavioral experiments, perfusion, sample collection, sectioning and staining in 4 doses experiment study. FS co-conceived the study, acquired funding support, manuscript revision and editing.

## Acknowledgements

The authors acknowledge the Mass Spectrometry Facility and Klaudia I. Hebert Kocurek, Senior Research Instrumentation Specialist, for proteomic assistance. The authors also acknowledge the NExT Department behavioral core facilities at the Texas A&M Vashisht College of Medicine. The schematic diagram was created using BioRender.com. This work was supported by the AHA Career Development Award (Award ID 25CDA1450365) to KKM and by RF1NS119872, the Woodnext Foundation and Cox Endowed Chair funds to FS.

## SUPPLEMENTARY FIGURES

**Fig. S1.**
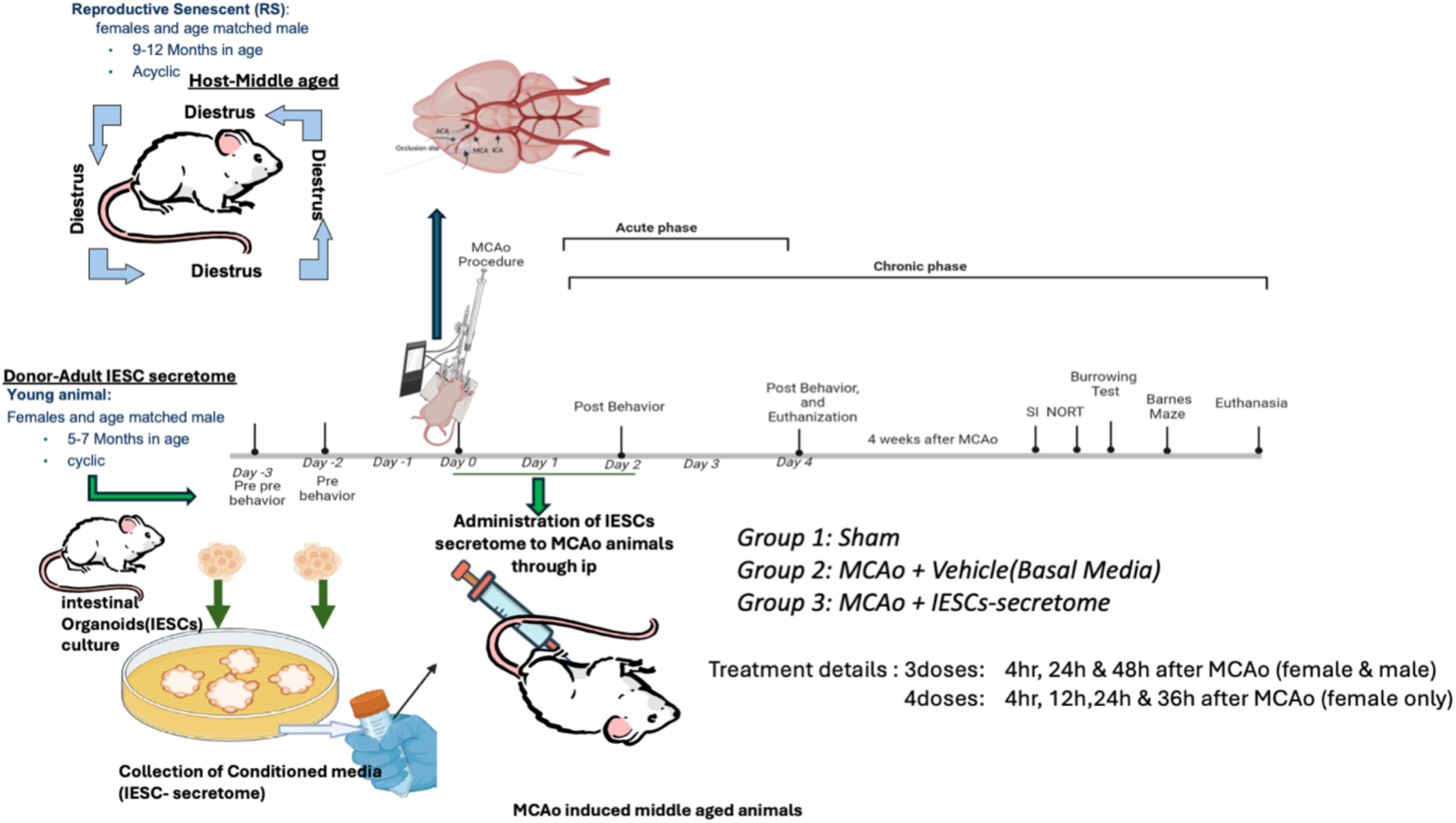
Experimental Design of the study. To investigate the therapeutic potential of intestinal epithelial stem cell-conditioned media (IESC-secretome) in stroke recovery, reproductively senescent (9–12 months old) female rats (acyclic) and age-matched males were subjected to transient middle cerebral artery occlusion (MCAo) or sham surgery. IESC-secretome was collected from organoid cultures from young male and female donor rats (5–7 months old).

**Fig. S2.**
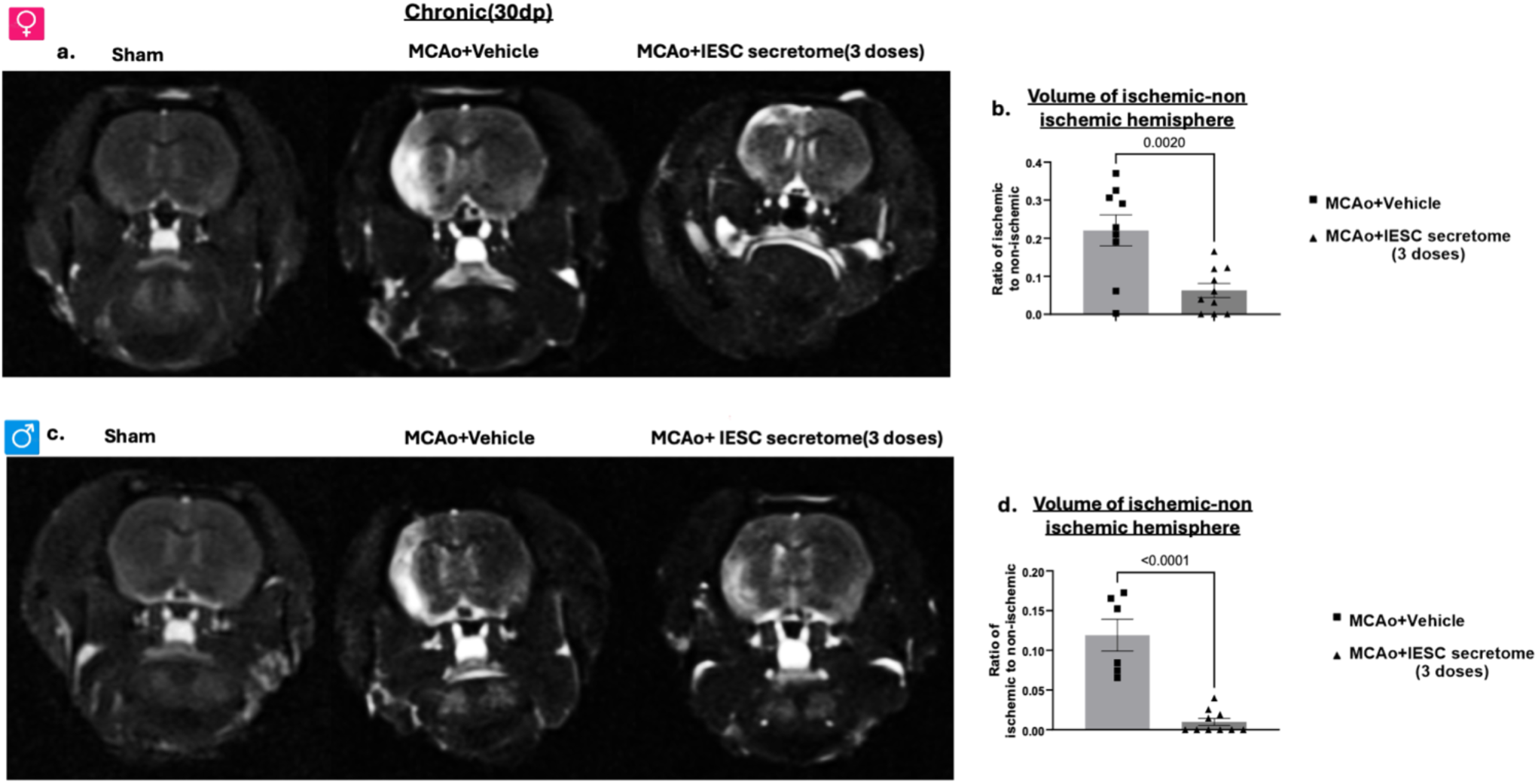
Effect of IESC-secretome treatment on T2 hyperintensity volume after stroke. Representative T2-weighted magnetic resonance imaging (MRI) axial slices from sham, MCAo + vehicle and MCAo + IESC-secretome–treated females (a) and males (c). Histograms depict non-normalized T2 hyperintensity volume normalized to the volume of the non-ischemic hemisphere (b,d). Data are presented as mean ± SEM; n=10/group. Data were analyzed by unpaired *t* test. Exact p values are shown on the graphs.

**Fig S3:**
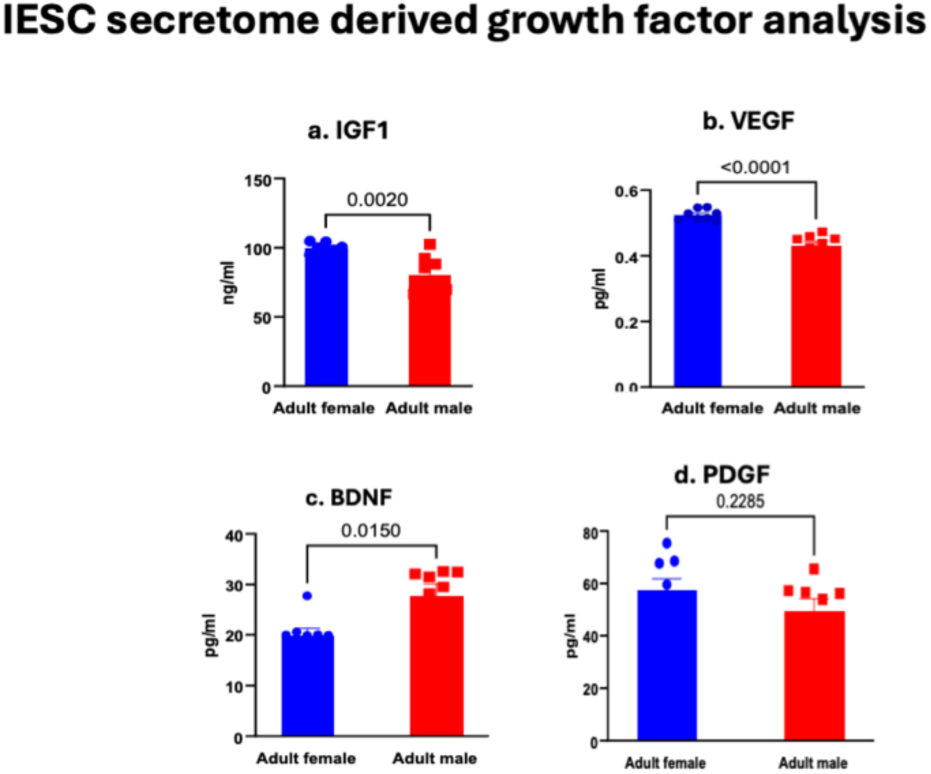
Sex-dependent differences in selected growth factors in IESC secretome. Low molecular weight growth factors (a) insulin-like growth factor 1 (IGF-1), (b) vascular endothelial growth factor (VEGF), (c) brain-derived neurotrophic factor (BDNF), and (d) platelet-derived growth factor (PDGF) were measured by ELISA in conditioned media from adult female and male IESC cultures. Graphs depict mean±SEM; n=9/group. Data were analyzed by t-test. Exact p values are shown on the graph.

**Fig. S4.**
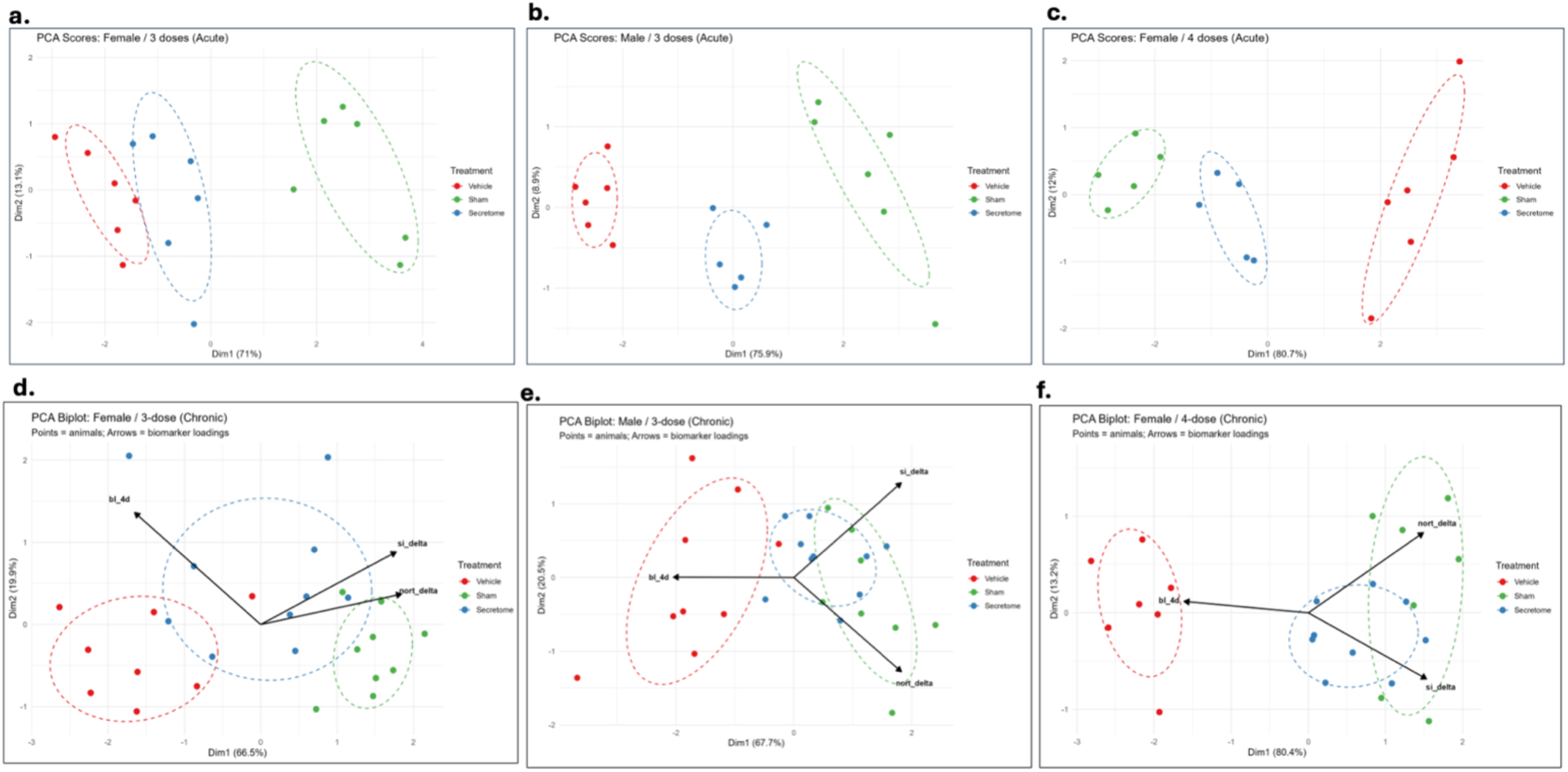
Principal component analysis of acute dataset: gut and immune biomarkers. Principal component analysis of gut/inflammatory biomarkers (acute) and behavioral outcomes (chronic). PCA was performed separately for each cohort using six gut/immune biomarkers (villus length-to-width ratio, goblet cell count, IL-17A, IL-6, IL-10, TNF-α) for the acute timepoint (a-c), and three behavioral measures (NORT_delta, SI_delta, and Day 4 Barnes maze latency) for the chronic timepoint (d-f). Points represent individual animals; dashed ellipses denote 68% confidence regions by treatment group (Vehicle, Sham, Secretome). (a) Acute, Female/3-dose (b) Acute, Male/3-dose (c) Acute, Female/4-dose (d) Chronic, Female/3-dose (e) Chronic, Male/3-dose (f) Chronic, Female/4-dose

